# The T-cell niche tunes immune function through modulation of the cytoskeleton and TCR-antigen forces

**DOI:** 10.1101/2024.01.31.578101

**Authors:** Anna V. Kellner, Rae Hunter, Priscilla Do, Joel Eggert, Maya Jaffe, Delaney K. Geitgey, Miyoung Lee, Jamie A. G. Hamilton, Anthony J. Ross, Raira S. Ank, Rachel L. Bender, Rong Ma, Christopher C. Porter, Erik C. Dreaden, Byron B. Au-Yeung, Karmella A. Haynes, Curtis J. Henry, Khalid Salaita

**Author notes:** Correspondence should be addressed to: K.S. and C.J.H.

## Abstract

Obesity is a major public health crisis given its rampant growth and association with an increased risk for cancer. Interestingly, patients with obesity tend to have an increased tumor burden and decreased T-cell function. It remains unclear how obesity compromises T-cell mediated immunity. To address this question, we modeled the adipocyte niche using the secretome released from adipocytes as well as the niche of stromal cells and investigated how these factors modulated T-cell function. We found that the secretomes altered antigen-specific T-cell receptor (TCR) triggering and activation. RNA-sequencing analysis identified thousands of gene targets modulated by the secretome including those associated with cytoskeletal regulation and actin polymerization. We next used molecular force probes to show that T-cells exposed to the adipocyte niche display dampened force transmission to the TCR-antigen complex and conversely, stromal cell secreted factors lead to significantly enhanced TCR forces. These results were then validated in diet-induced obese mice. Importantly, secretome-mediated TCR force modulation mirrored the changes in T-cell functional responses in human T-cells using the FDA-approved immunotherapy, blinatumomab. Thus, this work shows that the adipocyte niche contributes to T-cell dysfunction through cytoskeletal modulation and reduces TCR triggering by dampening TCR forces consistent with the mechanosensor model of T-cell activation.

## Introduction

The tissue microenvironment is a potent modulator of immune responses. T-cell function is regulated by extrinsic factors including inflammation, nutrient availability, and extracellular metabolites^1^. The loss of homeostasis, or the deregulation of these mediators can suppress immunity and promote immunological-driven pathologies^2^. For example, the progression of obesity is characterized by chronic inflammation and altered tissue metabolites^3,4^, which suppress the function of murine and human T-cells^5–8^. Obesity-induced T-cell immunosuppression is emerging as a risk factor for developing cancers of varying etiologies^9–12^, and more recently, in the context of solid tumors, has been shown to reduce the efficacy of some cancer therapies^13–15^. Currently, it is unknown how an obese microenvironment impacts immunotherapies targeting hematological malignancies such as chimeric antigen receptor (CAR) T-cells and bispecific T-cell engagers (BiTEs), which are FDA-approved treatments for patients with relapsed or refractory B-cell acute lymphoblastic leukemia (B-ALL), diffuse large B-cell lymphoma (DLBCL), multiple myeloma, and mantle cell lymphoma^14,16^ that rely on intrinsic cell function to eliminate cancerous cells^17–21^.

A fundamental question that relates to obesity-induced alterations in T-cell function is how the local microenvironment modulates basal signaling and antigen-triggered responses. Past literature indicates that high levels of leptin in the obese microenvironment increases PD1 receptor expression and T-cell exhaustion^12^, which could be detrimental to CAR-T-cell-based and BiTE immunotherapies. Additionally, increased fatty acid uptake in the tumor microenvironment leads to metabolic reprogramming of tumor-infiltrating T-cells and decreased T-cell cytotoxicity^11^. Lastly, it has been demonstrated that the stiffness of cancer cells *in vitro* dictates the T-cell response^22^. Despite the extensive documentation of obesity-induced T-cell dysfunction, the mechanisms underlying this association are still not fully understood.

One well-established mechanism of modulating T-cell responses involves the role of the T-cell cytoskeleton in tuning biophysical aspects of TCR triggering^23^. For example, it is established that the actin cytoskeleton couples to the TCR to mechanically regulate TCR triggering, immunological synapse formation, and target cell killing^23^. Additionally, cytoskeletal inhibitors are known to dampen TCR mechanical force generation, and therefore TCR triggering and T-cell activation^24^. Previously, our lab and others have shown that the T-cell cytoskeleton transmits 10-20 piconewton (pN)-level forces transmitted to the TCR-peptide major histocompatibility complex (pMHC) during antigen recognition^25–27^, which boosts antigen discrimination and enhances T-cell signaling by potentially altering receptor conformation and prolonging bond lifetime^24–31^. These forces are generated by actin polymerization and are transmitted to the TCR with the help of adapter proteins including pCasL, SLP-76, Fyb, and Nck, which are linked to actin polymerization proteins including WASP and Arp2/3^32,33^. Additionally, proper cytoskeletal function and actin retrograde flow are required for the formation of a functional immunological synapse between the T-cell and target cell^34–39^. Lastly, Huse and colleagues have shown that actin-mediated forces transmitted to the TCR and LFA-1 aid in target cell killing^40–42^. Accordingly, we sought to define how soluble factors in the microenvironment impact the T-cell cytoskeleton, TCR forces, and T-cell function.

In this work, we demonstrate that the secretomes of bone marrow stromal cells and adipocytes differentially impact T-cell function and the efficacy of blinatumomab, a BiTE immunotherapy commonly used to treat B-ALL. We first show that bone marrow stromal cell-conditioned media (SCM) and adipocyte-conditioned media (ACM) differentially alter T-cell activation. Functionally, SCM increased cytokine production (interferon-γ (IFN-γ)) and tumor necrosis factor-α (TNF-α)) and the expression of cytolytic mediators (perforin and GranzymeB (GzmB)). In contrast, ACM suppressed each of these parameters in T-cells as well as TCR signal transduction (upregulation of Nur77) and the surface expression of T-cell activation markers (CD25 and CD69). RNA sequencing of T-cells cultured in the secretomes of bone marrow stromal cells and adipocytes revealed rapid and distinctive gene expression changes in naïve T-cells. Notably, we observed that SCM induced high expression of critical adaptors or mediators that link actin polymerization to TCR signal transduction and immunological synapse formation, specifically *Lcp2*/SLP-76^32^. In contrast, ACM-mediated suppression of T-cell function occurred rapidly at the gene expression level in naïve T-cells with genes encoding TCR machinery and actin polymerization being significantly suppressed relative to levels observed in naïve T-cells cultured in unconditioned media or SCM. Next, we validated transcriptional differences in cytoskeletal genes by showing that SCM and ACM differentially alter total and filamentous actin (F-actin), as well as WAVE1 levels in the T-cell cytoskeleton at the protein level, with SCM leading to increased actin levels and ACM leading to actin and WAVE1 suppression. Due to the important role of the T-cell cytoskeleton in generating TCR-pMHC forces, we then employ DNA-based molecular force sensors to measure TCR forces in cells treated with conditioned media. We demonstrate that the microenvironment tunes TCR mechanical forces during initial antigen recognition, which validates the link between microenvironment-mediated alterations in the cytoskeleton and the biomechanics of the TCR. We then use a murine diet-induced obesity (DIO) model to demonstrate that T-cells isolated from the spleens of obese mice have significantly decreased cytokine secretion, TCR forces, and WAVE1 expression, showing that ACM mimics an obese *in vivo* environment. Lastly, using the BiTE, blinatumomab, which is an FDA-approved immunotherapy for B-ALL, we used a DNA-based molecular force sensor to show that the soluble microenvironment regulates mechanical forces transmitted through blinatumomab, as well as immunological synapse formation and target cell killing in co-culture assays of human T-cells with human B-ALL cells. Overall, our study demonstrates a novel microenvironmental mechanism regulating T-cell function and the efficacy of immunotherapies mediated by microenvironmental tuning of TCR mechanical forces.

## Results

### The soluble microenvironment modulates antigen-specific T-cell activation

As a model of T-cell activation, we utilized transgenic OT-1 mouse T-cells that specifically recognize the ovalbumin peptide (SIINFEKL) presented by H-2K^b^ ^43^. To mimic different microenvironments, we exposed T-cells to harvested conditioned media from bone marrow stromal cells and differentiated adipocytes (**Fig. 1a**). The procedure for generating conditioned media is shown in **Extended Data Fig. 1** and is previously described^44,45^. Our previous work has shown that adipocyte-secreted factors contain high levels of pro-inflammatory cytokines and chemokines including TNF-α and IL-6, which are indicators of chronic inflammation found in obesity^45^, and others have shown that soluble factors secreted by stromal cells enhance the immune response^46^. **SI Table 1** shows cytokine and chemokine expression levels in the conditioned media used in this work.

**Fig.1.**
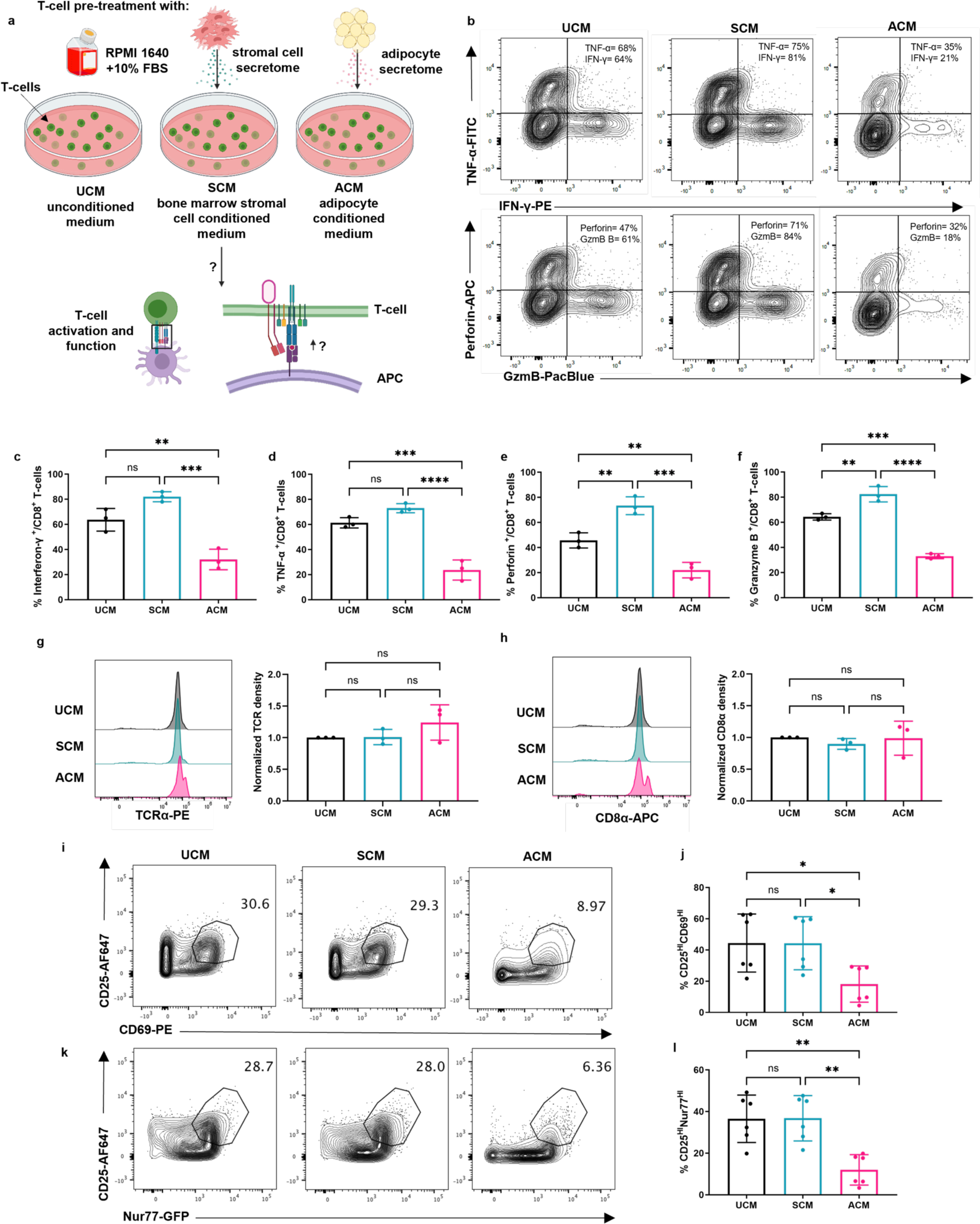
The soluble microenvironment modulates antigen-specific OT-1 T-cell activation. **a**, Schematic illustration of OT-1 T-cell activation in unconditioned medium (UCM), bone marrow stromal cell conditioned media (SCM), or adipocyte conditioned medium (ACM). **b,** Representative two-dimensional contour plots of tumor necrosis factor-α (TNF-α) and interferon-γ (IFN-γ) (top), or perforin and granzyme B (GzmB) (bottom). **c**, Percent of CD8+ OT-1 T-cells expressing IFN-γ after 72 hr activation. UCM v. SCM *P* = 0.0528, UCM v. ACM \*\**P* = 0.0047, SCM v. ACM \*\*\**P* = 0.0004 **d**, Percent of CD8+ OT-1 T-cells expressing TNF-α after 72 hr activation. UCM v. SCM *P* = 0.0982, UCM v. ACM \*\*\**P* = 0.0005, SCM v. ACM \*\*\*\**P* <0.0001. **e**, Percent of CD8+ OT-1 - cells expressing perforin after 72 hr activation. UCM v. SCM \*\**P* = 0.0047, UCM v. ACM \*\**P*= 0.0100, SCM v. ACM \*\*\**P* = 0.0002. **f**, Percent of CD8+ OT-1 T-cells expressing GzmB after 72 hr activation. **g**, TCR and **h**, CD8α expression of OT-1 T-cells after a 24 hr treatment with UCM, SCM or ACM. Data represents the TCR or CD8α geometric MFI of gated singlets normalized to UCM for each biological replicate. TCR: UCM v. SCM *P* = 0.9976, UCM v. ACM *P* = 0.2893, SCM v. ACM *P* = 0.3134, CD8α: UCM v. SCM *P* = 0.7383, UCM v. ACM *P* = 0.9964, SCM v. ACM *P* = 0.7830 **i**, Two-dimensional contour plots of CD25 and CD69 after a 24 h treatment with UCM, SCM, or ACM and a 2 hr stimulation with OVA peptide-pulsed splenocytes. Gate represents percent CD25-high and CD69-high OT-1 T cells. **j**, Quantification of percent CD25-high and CD69-high OT-1 T cells. UCM v. SCM *P* >0.9999, UCM v. ACM \**P* = 0.0315, SCM v. ACM \**P* = 0.0323. **k**, Two-dimensional scatter of CD25 and Nur77 after a 24 hr treatment with UCM, SCM, or ACM and a 2 hr stimulation with OVA-peptide-pulsed splenocytes. Gate represents percent CD25-high and Nur77-high OT-1 T cells. **l**, Quantification of percent CD25-high and Nur77-high OT-1 T cells. UCM v. SCM *P* = 0.9991, UCM v. ACM \*\**P* = 0.0020, SCM v. ACM \*\**P* = 0.0018. Graphs represent means ± SD. All data points represent independent biological replicates. All statistics were calculated using a one-way ANOVA with a Tukey’s multiple comparisons test. *n* = 3-6 independent animals.

To determine the effect of the local microenvironment on antigen-specific T-cell activation, OT-1 T-cells were cultured with OVA-peptide (SIINFEKL) loaded splenocytes in unconditioned medium (UCM), stromal cell-conditioned medium (SCM) or adipocyte-conditioned medium (ACM) for 3 days before assessing T-cell effector functions (**Fig. 1a**). Relative to responses observed in UCM, OT-1 T-cells stimulated in ACM exhibited a significant decrease in IFN-γ, TNF-α, perforin, and GzmB production by 49±9%, 61±7%, 52±12%, and 49±5%, respectively (**Fig. 1b-f**). In contrast, T-cells stimulated in SCM significantly increased their production of cytolytic mediators by 61±12% (perforin), and 27±5% (GzmB) compared to those stimulated in UCM (**Fig. 1b-f**). Overall, these data demonstrate that the secretome of bone marrow stromal cells augments T-cell function; whereas the adipocyte secretome is suppressive to antigen-specific T-cells, which adds to the growing body of literature of the immunoregulatory properties of both cell types.

### The soluble microenvironment regulates T-cell receptor signaling

Based on these observations, we next wanted to validate whether the conditioned media also modulated TCR signaling at earlier time points. We first assessed the impact of UCM, SCM, and ACM on the phenotype of naïve OT-1 T-cells, with a focus on the surface expression of the TCR and CD8 co-receptor. To this end, OT-1 T-cells were cultured for 24 hours in UCM, SCM or ACM (without stimulation). We found that T-cells did not exhibit significant alterations in their surface expression of the TCR nor the CD8α co-receptor, although a trend towards increased surface expression was noted in ACM (**Fig. 1g-h**). We next assessed if TCR signaling was differentially impacted by the bone marrow stromal cell and adipocyte secretomes given their impact on T-cell function. In these experiments, we used Nur77-GFP-expressing OT-1 T-cells^47^ given that upregulation of this transcription factor is an early response specific to TCR triggering^47–49^ and is a critical regulator of T-cell metabolism by controlling glycolysis and oxidative phosphorylation during T-cell activation^50^.

To assess differences in TCR signaling, T-cells were treated for 24 hours with UCM, SCM, or ACM prior to a 2-hour stimulation with OVA peptide-pulsed antigen presenting cells. After the 2-hour incubation period, we measured the intracellular upregulation of Nur77-GFP, as well as the surface expression of CD69 and CD25 given that these receptors are rapidly upregulated within a few hours of TCR stimulation^48,51,52^. Relative to UCM, T-cells treated with ACM had a 26±21% decrease in the number of CD25^HI^CD69^HI^ cells as well as a 59±16% decrease in the number of CD25^HI^Nur77-GFP^HI^ cells (**Fig. 1i-l**); whereas similar responses in these markers of TCR stimulation were observed in T-cells cultured in UCM compared to SCM (**Fig. 1i-l**). In all, these data demonstrate that the bone marrow stromal cell and adipocyte secretomes differentially tune the effector capacity of naïve T-cells prior to antigenic exposure by modulating TCR signaling properties.

### The soluble microenvironment alters the transcriptome of naïve CD3+ T-cells

To gain a comprehensive understanding of how bone marrow stromal cells and adipocyte secretomes differentially prime naïve T-cells, we performed RNA-sequencing analysis on naïve murine CD3+ polyclonal T-cells after a 4-hour culture in UCM, SCM or ACM. Note that live/dead staining via flow cytometry showed no significant changes in T-cell viability for all three conditions tested (**Extended Data Fig. 2**).Principal component analysis (PCA) confirmed strong reproducibility across replicates and revealed that early transcriptional changes in naïve CD3+ T-cells were largely influenced by the soluble microenvironment, with ACM-induced changes showing the largest divergence relative to gene expression profiles observed in T-cells cultured in UCM or SCM (**Fig. 2a**). In addition to inducing the greatest transcriptional changes, ACM induced a greater number of differentially expressed genes in naïve T-cells relative to those observed in T-cells cultured in SCM (**Fig. 2b-c**). To this point, 450 genes were significantly upregulated, and 698 genes were significantly downregulated in naïve T-cells cultured in ACM relative to UCM; whereas 97 genes were upregulated and 79 genes were downregulated in naïve T-cells cultured in SCM relative to UCM (**Fig. 2c**). Of these, 32.8% (415 genes) were uniquely upregulated in naïve T-cells cultured in ACM relative to UCM and 53.3% (675 genes) were downregulated only upon exposure to the adipocyte secretome (**Fig. 2c**). In contrast, 4.8% (61 genes) and 4.5% (57 genes) were uniquely up- and downregulated, respectively, in naïve T-cells cultured in SCM relative to UCM (**Fig. 2c**).

**Fig. 2.**
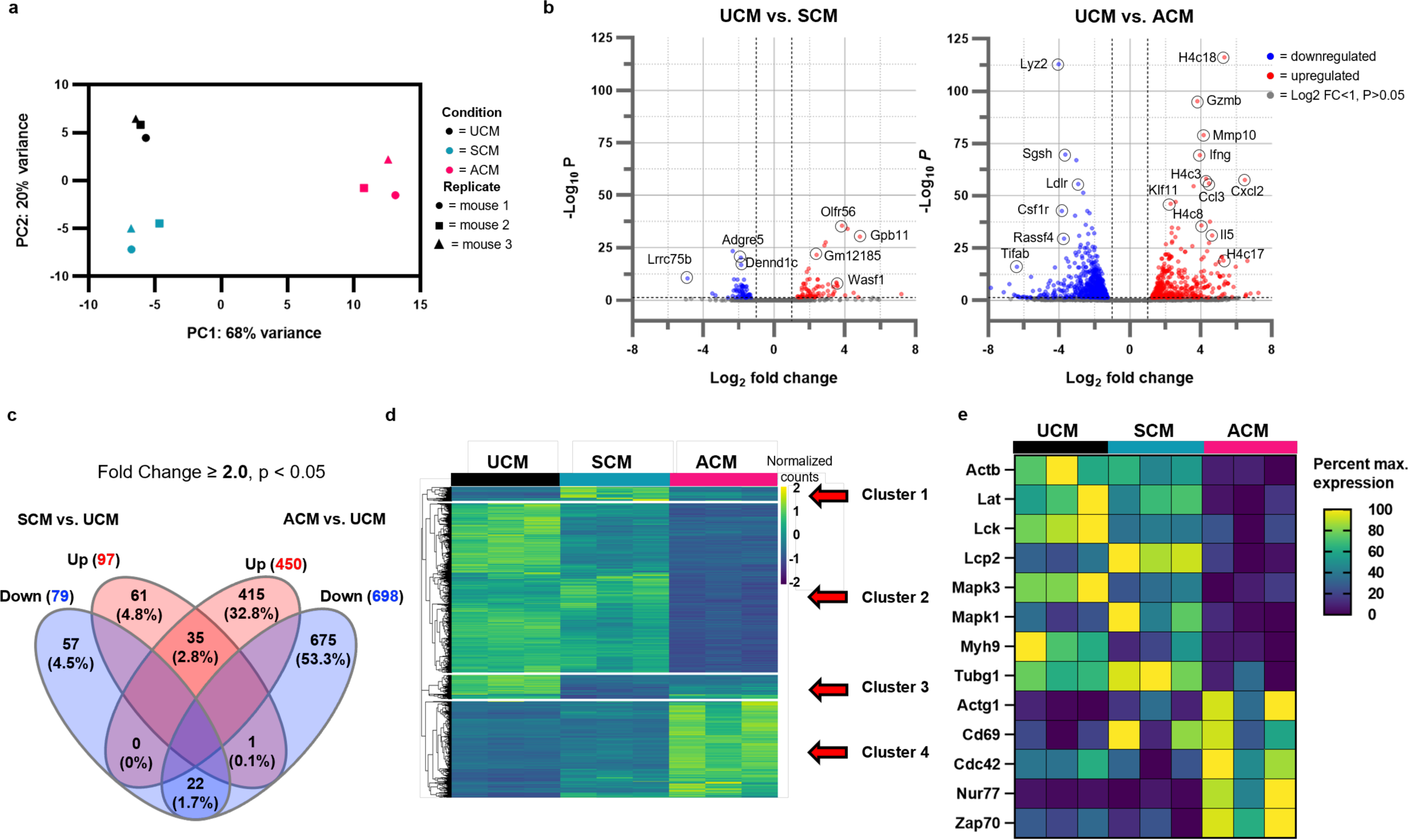
The soluble microenvironment alters gene expression profiles in naïve CD3^+^ T-cells. **a**, CD3+ T-cells were isolated from 3 separate C57BL/6 mice and treated for 4 hr with UCM, SCM, or ACM to observe early changes in gene expression via RNA-sequencing. PCA plot shows reproducibility between replicates. **b**, Volcano plots showing genes in SCM- or ACM-treated cells with a >2-fold change in expression compared to UCM. **c**, Venn Diagram showing overlap between genes that are upregulated or downregulated in SCM- and ACM-treated cells. **d**, Hierarchical clustering of 17,486 genes with an expression signal greater than zero in any condition. Cluster 1 = genes upregulated in SCM, Cluster 2 = genes downregulated in ACM, Cluster 3 = genes with a similar loss of expression in SCM and ACM, Cluster 4 = genes with increased expression in ACM. **e**, T-cell activation-specific targeted gene analysis. *n* = 3 independent animals.

Hierarchical clustering analysis further highlighted the unique transcriptional profiles induced by bone marrow stromal cell and adipocyte secretomes (**Fig. 2d**). From this analysis, gene expression profiles in naïve T-cells binned into four clusters when cultured in UCM, SCM, or ACM. Cluster 1 represents genes that were significantly increased in naïve T cells cultured in SCM relative to UCM and ACM. Of the top 40 uniquely upregulated genes in cluster 1, notable targets included genes involved in the cellular response to cytokine, actin cytoskeletal regulation, and T-cell activation (**Extended Data Fig. 3a**). Cluster 2 genes were significantly downregulated in ACM compared to UCM and SCM. Among these genes within cluster 2, we identified several transcripts involved in the response to external biotic stimuli, signal transduction, and cell adhesion (**Extended Data Fig. 3b**). Cluster 3 genes had a similar loss of expression in ACM and SCM compared to UCM, and therefore are not biologically interesting to this study because they provide no insight into what is happening uniquely in SCM or ACM. Lastly, cluster 4 represents genes that have an enhanced expression in ACM compared to UCM and SCM. Interestingly, a subset of upregulated cluster 4 genes were found to be involved in cytokine and chemokine activity, nucleosome assembly, apoptosis, and lipid metabolism (**Extended Data Fig 3c).** Interestingly, we noted an upregulation of genes involved in the type 2 inflammatory response (*TSLP*, *IL4*, *IL5*, and *IL13*), indicating a Th2 response ^53^ in T-cells treated with ACM. Previous work has shown that murine CD4+ T-cells gain a Th2 phenotype upon adoptive transfer into an obese mouse^54,55^.

We next identified a subset of 13 altered genes of interest that are sensitive to ACM or SCM treatment and are also involved in TCR signal transduction (*Lat*, *Lck*, *Lcp-2*, *Mapk1*, *Mapk3*, *Zap70*) ^56^, T-cell activation (*Nur77*, *CD69*) ^48^, or cytoskeletal activity (*Actb*, *Myh9*, *Tubg1*, *Cdc42*) ^57,58^ (**Fig. 2e**). Notably, *Actb*, *Lat*, *Lck*, *Lcp2*, *Mapk3*, *Mapk1*, *Myh9*, and *Tubg1* were lower in ACM-treated cells compared to UCM or SCM-treated cells (**Fig. 2e**). Downregulation of these genes would negatively impact proximal TCR signal transduction (*Lat*, *Lck, Mapk3*, *Mapk1*) ^59^ and immunological synapse formation (*Actb*, *Myh9*, *Tubg1*) ^60^, which hint at possible mechanisms underlying adipocyte-mediated suppression of T-cell function. Interestingly, we observed increased gene expression levels of *Lcp2* and *Mapk1*, the genes that encode SLP76 and ERK2, respectively, in naïve T-cells cultured in SCM (**Fig. 2e**). Notably, *Lcp2*/SLP76 is a master regulator of T-cell function, where it serves as a critical adaptor for TCR proximal downstream signaling molecules^59^ and a nucleation hub for actin polymerization^32,61^. An upregulation of *Lcp2* in SCM-cultured naïve T-cells could potentially explain the enhanced TCR signaling (**Fig. 1**) and augmented effector functions (**Fig. 1**) observed in T-cells in this condition. In addition, ERK2 phosphorylation occurs soon after TCR triggering and is a crucial step in the signal transduction cascade that leads to T-cell activation^62^. An upregulation of *Mapk1*/ERK2 provides an additional explanation for the enhanced TCR signaling seen in **Fig. 1**. Genes that were upregulated in naïve T-cells exposed to the adipocyte secretome include *Actg1*, *Cd69*, *Cdc42*, *Nur77*, and *Zap70* (**Fig. 2e**). These results indicate that these genes can be induced in an antigen-independent manner and may be upregulated in response to inflammatory cytokines or as a compensatory mechanism to overcome adipocyte-mediated T-cell suppression.

Given the number of differentially expressed genes associated with TCR signal transduction, actin nucleation, and the formation of the immunological synapse, we decided to further investigate how the bone marrow stromal cell and adipocyte secretomes modulated β-actin and cytoskeletal dynamics in naïve T-cells.

### The soluble microenvironment changes total actin and filamentous actin levels in OT-1 T-cells

To gain a better understanding of how bone marrow stromal cells and adipocytes impact cytoskeletal function, we identified differentially expressed genes involved in the regulation of the actin cytoskeleton (**Fig. 3a-b**). Wiskott-Aldrich syndrome protein family member 1 is a protein coded by the *Wasf1* (*Wave1*) gene^63^ and we observed a significant increase in the expression of this gene in SCM-cultured T-cells (**Fig. 3a-b**). *Wasf1* plays a critical role in the WASP-family verprolin homologous protein (WAVE1) regulatory complex which enhances actin polymerization through its association with Rac and the actin nucleation core Arp2/3 complex^64^. This protein also enhances cytoskeletal membrane ruffling, thereby promoting a motile cell surface^65,66^. Notably, this was the only actin-related gene that was increased in naïve T-cells cultured in SCM relative to other conditions tested (**Fig. 3a-b**). In contrast, ACM upregulated genes that inhibit actin cytoskeletal function (*Ppp1r12a*, *Raf1*, and *Pak1*) and downregulated genes that enhance actin polymerization, adhesion, and cytoskeletal activity (*Mylk3*, *Rac2*, *Itga4*, *Actn1*, *Pik3r2*, *Itga7*, *Arghef4*, *Lpar5*, *Pip4k2b*, *Pfn2*, *Iqgap3*, *Itga1*, *Pdgfb*, and *Itga5*) (**Fig. 3a-b**).

**Fig. 3.**
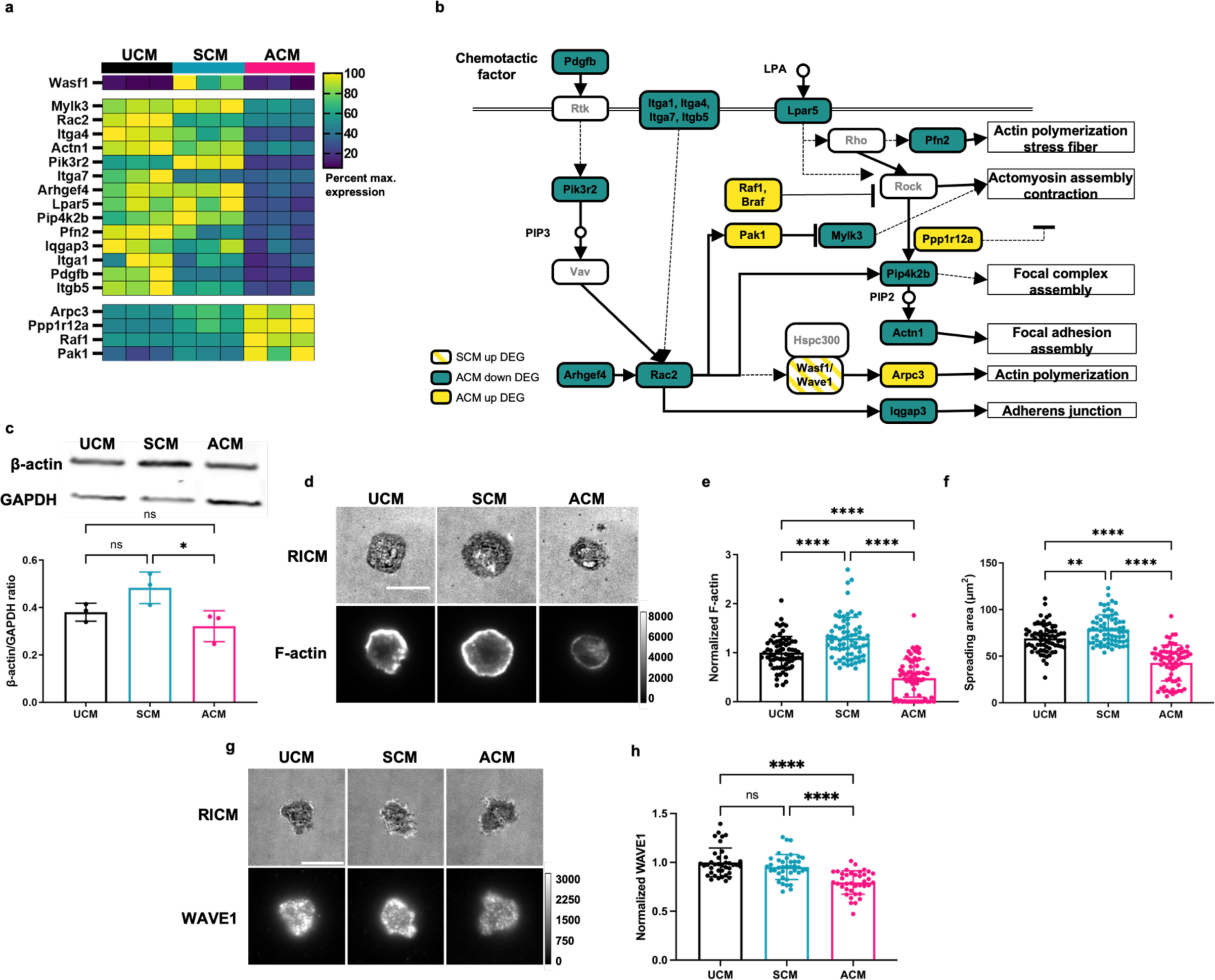
Soluble factors secreted by bone marrow stromal cells and adipocytes differentially regulate actin homeostasis and polymerization in CD3+ lymphocytes. **a**, Expression differences in genes involved in actin cytoskeleton regulation (KEGG hsa04810). **b**, Map highlighting gene expression differences in the actin cytoskeleton regulation pathway (modified from KEGG hsa04810). **c**, Western blot quantifying β-actin protein expression in OT-1 T-cells treated for 24 hr in UCM, SCM, or ACM. UCM v. SCM *P* = 0.1542, UCM v. ACM *P* = 0.4708, SCM v. ACM \**P* = 0.0325. Data points represent individual biological replicates. **d**, Representative images of actin polymerization (F-actin) measured with SirActin in OT-1 T-cells treated for 24 hr in UCM, SCM, or ACM and **e**, quantification of F-actin fluorescence intensity. Individual data points were normalized to the mean fluorescence intensity of UCM in each biological replicate. \*\*\*\**P* <0.0001. **f**, Cell adhesion of OT-1 T-cells treated with UCM, SCM, or ACM for 24 hr measured via cell spreading area. \*\**P = 0.0024, ****P <* 0.001. **g**, Representative images of WAVE1 measured with immunofluorescence in OT-1 T-cells treated for 24 h in UCM, SCM, or ACM and **h**, quantification of WAVE1 fluorescence intensity. Individual data points were normalized to the mean fluorescence intensity of UCM in each biological replicate. ****P<0.0001. Scale bars = 10 μm. Data points represent individual cells in **e-f, h**. Graphs represent means ± SD. All statistics were calculated using a one-way ANOVA with a Tukey’s multiple comparisons test. *n* = 2-4 independent animals.

To validate gene expression changes at the protein level in naïve T-cells in different microenvironments, we measured total β-actin levels in OT-1 T-cells treated with UCM, SCM, or ACM for 24 hours. We observed a significant decrease in the β-actin/GAPDH ratio in cells treated with ACM compared to SCM with no significant changes in GAPDH levels between conditions (**Fig. 3c** and **WB Source Data**) Next, we measured filamentous actin (F-actin) levels, cell adhesion, and WAVE1 protein expression to corroborate ACM-mediated changes in the cytoskeleton in these cells. For these experiments, we treated OT-1 T-cells for 24 hours with UCM, SCM, or ACM prior to seeding on anti-CD3ε-coated glass for 30 minutes. Cells were then fixed, permeabilized, and stained with SirActin to measure F-actin levels or anti-WAVE1 to measure WAVE1 expression via fluorescence microscopy. Cell spreading area was also measured for these cells using RICM. Compared to UCM, OT-1 T-cells treated with ACM had a 52±6% decrease in F-actin levels as well as a 38±4% decrease in spreading area, and a 21±3% decrease in WAVE1 levels (**Fig. 3d-h**). In contrast, OT-1 T-cells treated with SCM had a 29±6% increase in F-actin levels as well as a 14±4% increase in spreading area, and no significant change in WAVE1 expression compared to cells treated with UCM (**Fig. 3d-h**). Additionally, we measured total cholesterol levels using Filipin III staining after treatment with UCM, SCM, or ACM, because previous work demonstrated that membrane cholesterol can impact membrane stiffness and TCR triggering, and therefore may be responsible for a change in T-cell function^67,68^. Compared to UCM, we observed a 20±8% and 24±8% increase in cholesterol content in ACM and SCM, respectively (**Extended Data Fig. 4a-b**). These results rule out cholesterol modulation as the sole mechanism of differential T-cell regulation in SCM and ACM.

Given that F-actin polymerization and WAVE1 activity are major mediators of TCR forces^25,69^ and we saw significant changes in F-actin and WAVE1 levels in SCM and ACM, we next sought to determine the effects of the adipocyte and bone marrow stromal cell secretomes on mechanical forces exerted by the TCR.

### The soluble microenvironment alters TCR biomechanics

Work by our group and others demonstrated that pN-level forces are an important component of TCR triggering upon antigen recognition^25–27,29,30^. Since ACM compromised T-cell function and suppressed TCR signal transduction, β-actin protein levels, and F-actin polymerization, we hypothesized that T-cell suppression is mediated by dampened TCR forces upon antigen recognition. Additionally, we expected SCM treatment to enhance TCR biomechanics due to enhanced T-cell signaling upon treatment with SCM (**Fig. 1**) and enhanced F-actin levels (**Fig. 3**).

We employed a type of molecular force sensor termed DNA-based tension probes to measure TCR forces during antigen recognition^25,70–72^ (**Extended Data Fig. 5** and **Table S2**). DNA-based tension probes consist of a DNA duplex that is anchored to a glass surface on one end and presents a ligand of interest to the T-cell on the other end. The DNA duplex has a hairpin that is flanked by a fluorophore-quencher pair, and the hairpin mechanically unfolds when the TCR exerts a threshold mechanical force on the ligand. The mechanical unfolding of the hairpin separates the fluorophore and quencher, causing a large increase in fluorescence intensity that can be mapped and measured in real-time by conventional fluorescence microscopy^25,70–72^. The force at which the hairpin mechanically unfolds, or the F_1/2_, is defined by the force at which the hairpin has a 50% probability of unfolding^73,74^. The F_1/2_ is tuned by altering the GC content of the hairpin and the size of the stem loop^73,74^. In the experiments described below, we used a hairpin with F_1/2_ = 4.7 pN to measure mechanical forces exerted by the TCR in different microenvironments. First, we performed a proof-of-concept experiment by culturing isolated polyclonal CD4+ and CD8+ murine T-cells in UCM, SCM, or ACM for 24 hours before seeding them on DNA-based tension probes presenting an anti-CD3ε antibody as the ligand (**Fig. 4a**). Relative to UCM, CD4+ and CD8+ T-cells cultured in ACM had a 41±4% and 34±4% decrease in 4.7 pN tension signal, respectively (**Fig. 4b-c**). In contrast, CD4+ and CD8+ cultured in SCM had a 278±14% and 416±23% increase in 4.7 pN tension signal, respectively (**Fig. 4d-e**). Next, we validated these responses in antigen-specific T-cells. OT-1 T-cells were cultured in each condition prior to seeding on DNA-based tension probes presenting the H2-K^b^ OVA 257-264 SIINFEKL epitope that is specifically recognized by the OT-1 TCR (**Fig. 4f**). Relative to UCM, OT1 T-cells cultured in ACM had a 37±6% decreased in TCR tension signal (**Fig. 4g**). In contrast, OT1 T-cells cultured in SCM had a 1240±84% increase in TCR tension signal (**Fig. 4h**). Additionally, OT-1 T-cells treated with SCM were able to exert forces of higher magnitudes (12 and 19 pN) compared to UCM (**Extended Data Fig. 6**). Previously, our lab has shown that OT-1 T-cells were unable to mechanically unfold 19 pN hairpin probes unless ICAM was present^25^, demonstrating that the soluble microenvironment significantly modulates the magnitude of TCR mechanics. Timelapse videos showed that OT-1 TCR forces were highly dynamic for all three environments (UCM, SCM, and ACM), but as quantified above, ACM forces were significantly dampened (**Supplementary movies 1-3**).

**Fig. 4.**
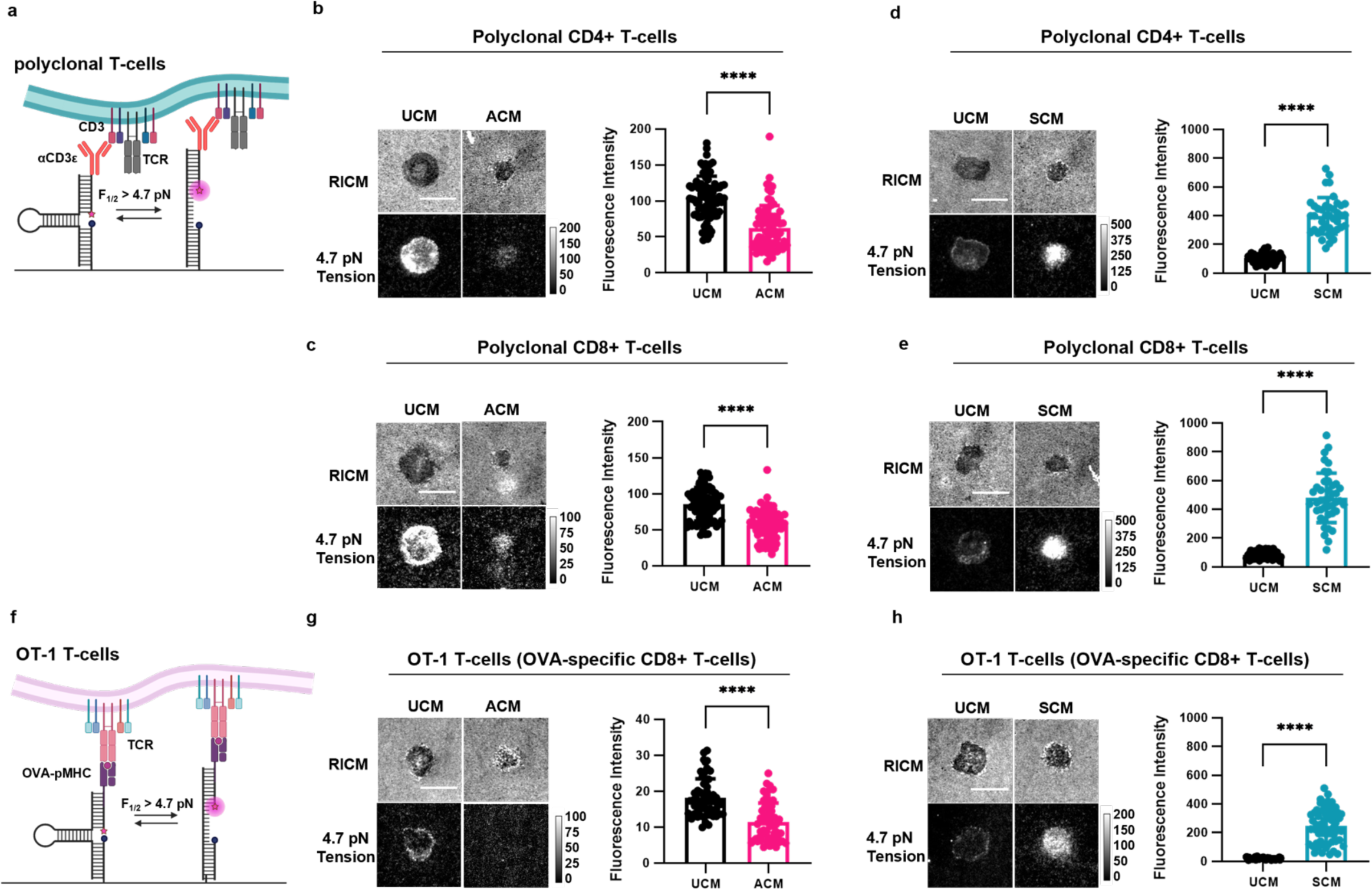
Soluble factors in the microenvironment modulate TCR biomechanics. **a**, Schematic of a DNA-based tension probe presenting α-CD3ε. **b**, Representative images and quantification of 4.7 pN tension signal exerted by polyclonal CD4+ and **c**, CD8+ T-cells treated with UCM or ACM for 24 hr. **d**, Representative images and quantification of 4.7 pN tension signal exerted by polyclonal CD4+ and **e**,CD8+ T-cells treated with UCM or SCM for 24 hr. **f,** Schematic of a DNA-based tension probe presenting the SIINFEKL OVA-pMHC to antigen-specific OT-1 T-cells. **g**, Representative images and quantification of 4.7 pN tension signal exerted by OT-1 T-cells treated with UCM or ACM for 24 hr and **h**, UCM or SCM for 24 hr. Scale bars = 10 μm. Graphs represent means ± SD and data points represent fluorescence intensity measurements of individual cells. Statistics were calculated using a two-tailed student’s t-test. \*\*\*\**P* < 0.0001, *n* = 2-3 independent animals.

### Diet-induced obesity dampens T-cell function, TCR forces, and cytoskeletal activity in a mouse model

We next tested whether diet induced obesity (DIO) can recapitulate the observed modulation in T-cell function and TCR forces mediated by ACM. To achieve this, we fed C57BL/6 mice high fat diet (HFD) or control diet (CD) for 3 months to induce obesity (**Fig. 5a**). Isolated T-cells from these mice were stimulated with anti-CD3/anti-CD28 coated beads for 72 hrs and then TNF-α, IFN-γ, GzmB, and Perforin levels were quantified using flow to infer the fraction of activated CD4+/CD8+ T-cells (**Fig. 5b-g**). Across all measured indicators of T-cell activation, we found a significant decrease in T-cells harvested from obese animals. This is consistent with our results showing that ACM dampens T-cell activation in OT-1 cells (**Fig. 1**). TCR force measurements using force probes presenting anti-CD3 demonstrated a 29±5% decrease in TCR forces (4.7 pN) for T-cells obtained from animals on HFD compared to the CD group (**Fig. 5h-i**). Interestingly, this observation matched our force measurements on polyclonal CD4+ and CD8+ T-cells treated with ACM and that showed a ∼30-40% decrease in tension signal (**Fig. 4b** and **4c**). Finally, we quantified WAVE1 expression and cell spreading area on anti-CD3 presenting surfaces and found a 27±6% and 25±5% reduction in WAVE1 expression and in cell spreading area, respectively (**Fig. 5j-l**). Taken together, DIO leads to functional alterations in T-cell activation, TCR forces, and T-cell cytoskeletal activity that matches that observed due to ACM treatment and suggests similar mechanisms of action.

**Fig.5.**
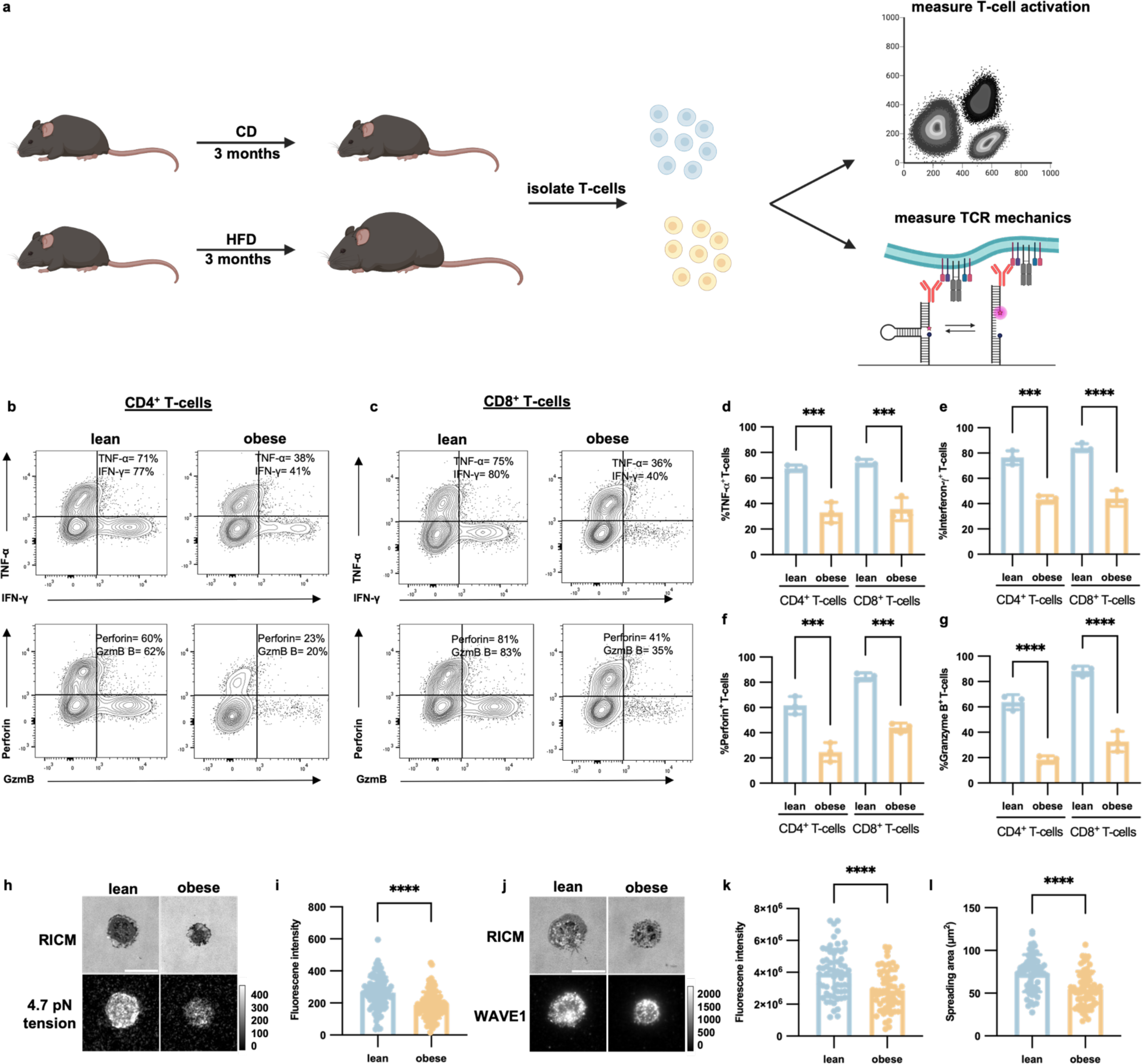
T cells from DIO mice have reduced cytokine production and TCR biomechanics. **a**, Schematic illustration of *ex vivo* T cell activation experiments performed on mice fed with a control diet (CD, lean) or high fat diet (HFD, obese) for 3 months**. b,** Representative two-dimensional contour plots of tumor necrosis factor-α (TNF-α) and interferon-γ (IFN-γ) (top), or perforin and granzyme B (GzmB) (bottom) for CD4+ and **c,** CD8+ T cells. **d**, Percent of T cells expressing TNF-⍺ (CD4+ ***P = 0.007, CD8+ ***P = 0.005) **e**, IFN-ɣ (***P = 0.0001, CD8+ P<0.0001) **f**, perforin (CD4+ ***P = 0.0002, CD8+ ***P = 0.0001), or **g**, granzyme B (****P<0.0001) after 72 h stimulation. Statistics were calculated using a one-way ANOVA with a Tukey’s multiple comparisons test. N = 3 mice per group. **h**, Representative images and **i**, quantification of 4.7 pN tension signal exerted by CD3+ T cells isolated from lean or obese mice (****P<0.0001). N = 3 mice from each group. **j**, Representative images and **k**, quantification of WAVE1 fluorescent intensity and **l**, cell spreading area of CD3+ T cells isolated from lean and obese mice (****P<0.0001). Statistics were calculated using an unpaired T test. N = 2 mice per group.

### T-cell transmitted forces to the FDA-approved immunotherapy, blinatumomab, are regulated by the soluble microenvironment

We next explored whether SCM and ACM modulate the magnitude of TCR forces transmitted through the BiTE, blinatumomab, which is an FDA-approved drug to treat B-cell lymphoblastic leukemia. Blinatumomab acts by bridging CD3+ T-cells with CD19+ B-ALL cells to induce B-ALL cytolysis^75^ (**Fig. 6a**). Here, we hypothesized that SCM and ACM modulation of T-cell function and TCR-antigen forces would be mirrored using BiTEs with human donor T-cells. To test this hypothesis, we designed a DNA-based tension probe that presents CD19 as its ligand. Next, we added blinatumomab to the CD19 probes to mimic the role of the BiTE in patients (**Fig. 6b** and **Extended Data Fig. 7**). We seeded CD3+ human donor T-cells on the blinatumomab-presenting probes (“blina-probes”) and amplified the signal using a DNA strand that is complementary to the stem loop region of the hairpins to lock the probes open after they are mechanically unfolded by the T-cell^76,77^ (**Fig. 6b**). Using the locking strand, we measured the accumulation of TCR tension signal generated by T-cells cultured in UCM, SCM, or ACM. (**Fig. 6c-d**). Compared to UCM, we observed a 24±8% increase in accumulated tension signal generated by T-cells exposed to SCM; whereas the ACM resulted in a 53±8% decrease in accumulated tension signal when human T-cells were added to blina-probes. To show that binding and tension were specific to blinatumomab, we seeded T-cells on probes presenting only CD19 and observed no binding or accumulation of tension signal (**Extended Data Fig. 8**). These results demonstrate that human TCR biomechanical forces are modulated by the soluble microenvironment and dictate the strength of the interaction with the FDA-approved drug blinatumomab.

**Fig. 6.**
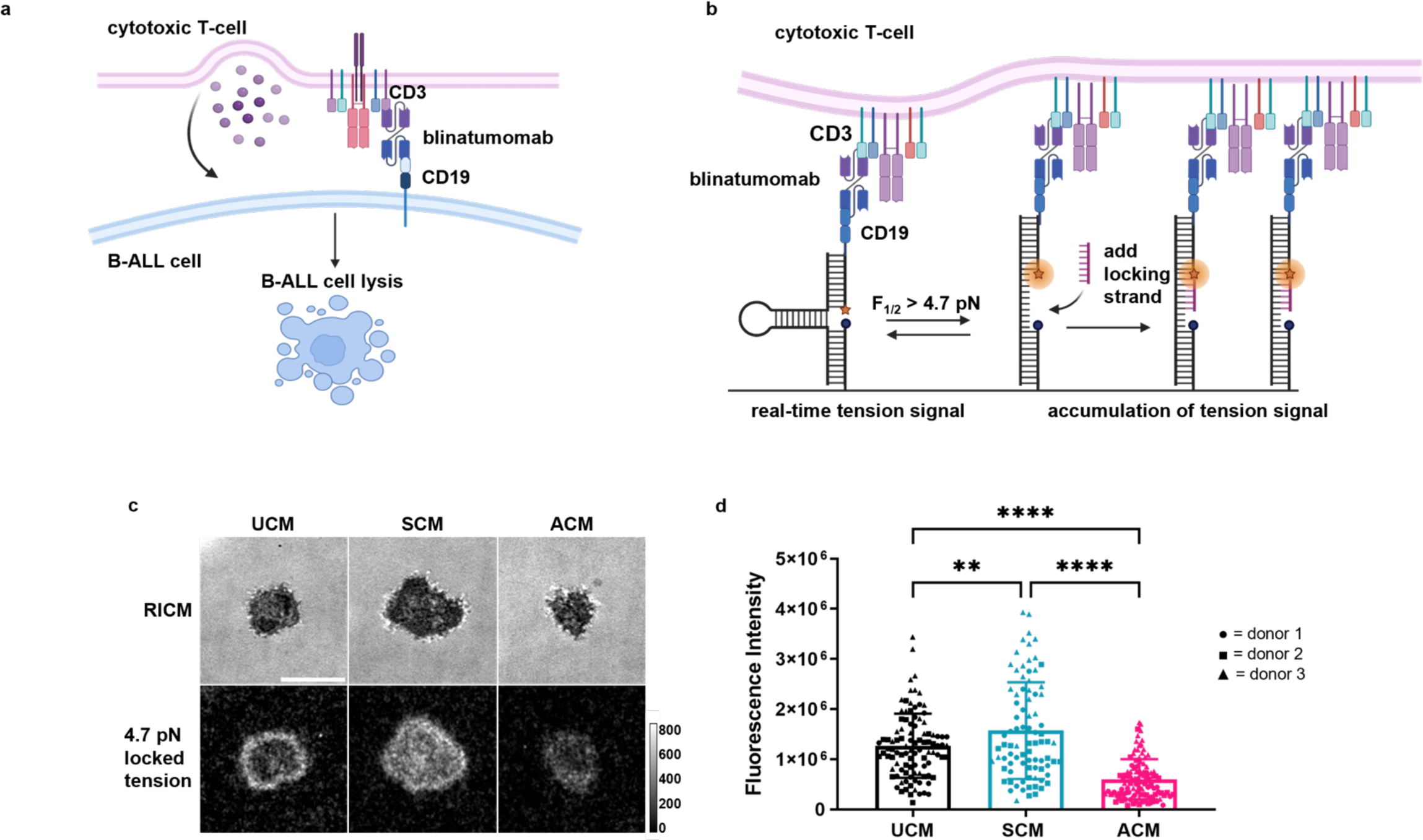
The biomechanics of the immunotherapy, blinatumomab, are regulated by the soluble microenvironment. **a**, Schematic of the mechanism of action of blinatumomab. Blinatumomab is a bispecific T-cell engager (BiTE) that bridges CD3+ T-cells with CD19+ B-cells to induce cytolysis. **b**, Schematic of the blinatumomab DNA-based tension probe design (“blina-probe”). Blinatumomab is added to CD19-presenting tension probes to bridge CD3+ T-cells with CD19-presenting probes. **c**, Representative images and **d**, quantification of locked 4.7 pN tension signal of donor T-cells on blina-probes after a 24 hr treatment with UCM, SCM, or ACM. Circles (●) represent cells from donor 1, squares (▪) represent cells from donor 2, triangles (▲) represent cells from donor 3. Scale bar = 10 μm. Graphs represent means ± SD and data points represent fluorescence intensity measurements of individual cells. Statistics were calculated using a one-way ANOVA with a Tukey’s multiple comparisons test. **P < 0.0091, ****P < 0.0001, *n* = 3 independent donors.

### The soluble microenvironment impacts immunological synapse formation and blinatumomab-mediated killing of malignant B-cells

Since mechanical forces are important to TCR triggering^25–27^, immunological synapse formation^37,38^, and cytotoxicity^40–42^, we wanted to determine if reduced mechanical forces through blinatumomab correlated with reduced T-cell function. We assessed immunological synapse formation between human CD3+ T-cells and human B-ALL cells by gating on concentrated positive actin signal within the synapse between one T-cell and one B-ALL cell using ImageStream flow cytometry (**Fig. 7a-c**). We measured frequency of immunological synapse formation in human donor T-cells treated with blinatumomab compared to a vehicle control (**Fig. 7a-c, Extended Data Fig. 9**). Using human T-cells from two donors, we saw similar levels of immunological synapse formation between UCM- and SCM-treated cells in the presence of blinatumomab, whereas ACM reduced blinatumomab-mediated immunological synapse formation (**Fig. 7a-c**). Next, we assessed human T-cell-mediated cytolysis of human B-ALL cells in the absence and presence of blinatumomab after 24 and 48 hours of co-culture. Compared to UCM, human T-cells exposed to the bone marrow stromal cell secretome had a 51±12% increase in human B-ALL cell lysis while those exposed to the adipocyte secretome exhibited a 51±12% decrease in human B-ALL cell lysis after 24 hours of co-culture (**Fig. 7d**). After 48 hours of co-culture, increased killing was observed for human T-cells cultured in UCM and SCM; whereas, human T-cells failed to kill target B-ALL cells after 2 days of co-culture with blinatumomab in the adipocyte secretome (**Fig. 7e**).

**Fig. 7.**
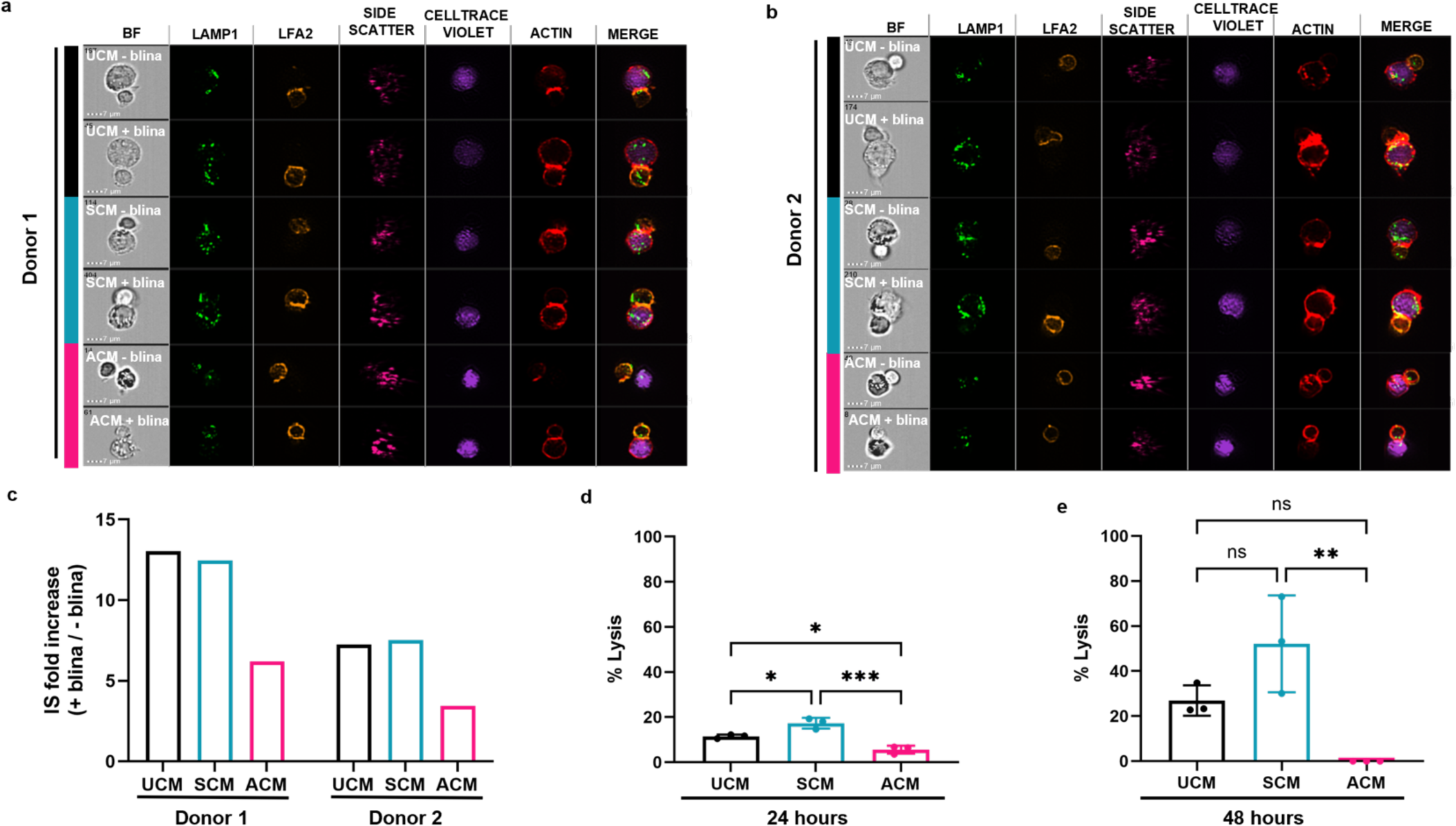
The soluble microenvironment impacts blinatumomab-mediated killing of malignant B-cells. **a-b**, Representative ImageStream flow cytometer images of blinatumomab-mediated immunological synapse formation after a 24 hr treatment with UCM, SCM or ACM. T-cells were stained with LAMP-1, LFA-2, and SirActin to show markers of immunological synapse formation. Nalm6 B-cells were stained with cell tracker violet. Scale bars = 7 μm. **c**, Fold-increase of immunological synapse formation in blinatumomab-treated samples compared to vehicle-treated samples. **d**, Blinatumomab-mediated cytolysis of Nalm6 B-cells after a 24 hr or **e**, 48 hr treatment with UCM, SCM, or ACM. %Lysis was calculated using 100 x (% blinatumomab-treated sample [Dead Nalm6] - %vehicle-treated sample [Dead Nalm6]). 24 hr: UCM v. SCM \**P* = 0.0151, UCM v. ACM \**P* = 0.0151, SCM v. ACM \*\*\**P* = 0.0004. 48 hr: UCM v. SCM *P* = 0.1201, UCM v. ACM *P* = 0.0971, SCM v. ACM \*\*\**P* = 0.0064. Graphs represent means ± SD. Data points represent individual donors. Statistics were calculated using a one-way ANOVA with a Tukey’s multiple comparisons test. *n* = 2-3 independent donors.

## Discussion

In these studies, we show that the soluble microenvironment dictates T-cell function, in part, by modulating the actin cytoskeleton and tuning TCR biomechanical forces (**Fig. 8**). Overall, our results show that soluble factors secreted by bone marrow stromal cells enhance TCR mechanical forces, which have been shown to promote T-cell activation and target cell killing^25,40^ (**Fig. 8a**). Additionally, soluble factors secreted by adipocytes weaken TCR biomechanical forces, which provides a novel suppressive mechanism explaining adipocyte-mediated T-cell dysfunction (**Fig. 8b**).

**Fig. 8.**
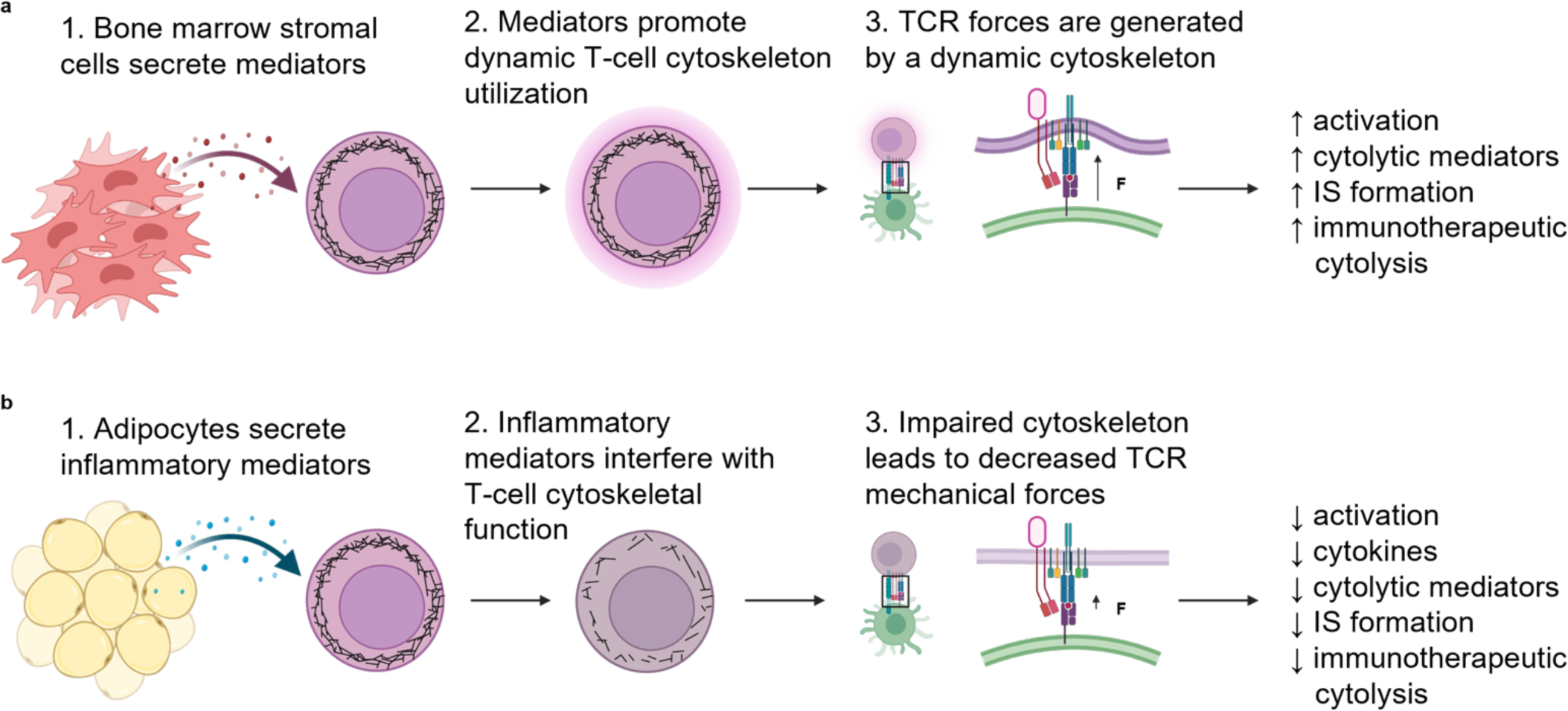
Proposed model showing how the soluble microenvironment may impact T-cell activity. **a,** Schematic illustration showing how the bone marrow stromal cell secretome may impact T-cell activation. Bone marrow stromal cells secrete soluble factors into the surrounding microenvironment. These soluble mediators promote optimal usage of the T-cell cytoskeleton, which leads to enhanced TCR forces and therefore increases T-cell activation, production of cytolytic mediators, immunological synapse formation, and immunotherapeutic-mediated cytolysis. **b**, Schematic illustration showing how the adipocyte secretome may impact T-cell activation. Adipocytes secrete inflammatory mediators into the surrounding microenvironment. These soluble mediators interfere with the T-cell cytoskeleton, which leads to dampened TCR forces and therefore decreases T-cell activation, production of cytokines and cytolytic mediators, immunological synapse formation, and immunotherapeutic-mediated cytolysis.

The actin cytoskeleton has been shown to be a master regulator of T-cell function. Previous work demonstrated that substrate stiffness and the mechanical environment led to differential expression of genes involved in T-cell activation and the cytokine response^78^, showing that T-cells use their cytoskeleton to sense and adapt to local surroundings. To explore this concept at a molecular level, a biomembrane force probe (BFP) was used to show that inhibiting actin polymerization by adding latrunculin to the local environment leads to a decrease in both TCR-pMHC binding affinity and on-rate^79^. In addition, traction force microscopy and DNA-based tension probe experiments showed that latrunculin suppresses T-cell force generation. ^76,80^ Our work expands upon these results by measuring changes in actin polymerization and forces directly exerted by the TCR, as well as T-cell activation markers and effector functions, after exposure to complex bone marrow stromal cell and adipocyte secretomes. Additionally, we have uncovered a set of cytoskeletal genes that are significantly downregulated upon ACM treatment of naïve T-cells which may be responsible for the changes in TCR mechanics. These genes include *Mylk3*, *Itga4*, *Actn1*, *Pik3r2*, *Itga7*, *Arghef4*, *Lpar5*, *Pip4k2b*, *Pfn2*, *Iqgap3*, *Itga1*, *Pdgfb*, and *Itga5*, and play a crucial role in actin polymerization, stress fiber formation, actomyosin assembly and contraction, focal adhesion formation, and adherens junction formation. In addition, we showed that *Wasf1 (Wave1)* is uniquely upregulated in T-cells exposed to the bone marrow stromal cell secretome, providing a potential mechanism for the increased actin polymerization and TCR mechanics seen in these cells. Taken together, these results demonstrate that the actin cytoskeleton and endogenous TCR forces are differentially affected by adipocytic and stromal cell microenvironments. Furthermore, we demonstrate that adipocyte-secreted factors may hinder T-cell function by interfering with cytoskeletal mechanics. While we focused this study on cytoskeletal-related genes, these genes only represent a small fraction of the modulated gene pool. Therefore, there are certainly additional mechanism for how SCM and ACM alter T-cell function that we have yet to explore including altered T-cell metabolism.

Recently, cellular biomechanics have become a topic of interest in the field of immuno-oncology, as T-cell mechanical forces have been shown to play an important role in cancer cell killing^40,41,81,82^. In support of this, previous work using atomic force microscopy (AFM) and magnetic tweezers has shown that cancer cells with a high metastatic potential tend to be mechanically softer than cancer cells with a low metastatic potential^83,84^, which provides a potential mechanism for immune evasion because T-cells cannot accumulate F-actin at the immunological synapse and exert strong mechanical forces on mechanically soft cells^22^. Our work expands upon these concepts by showing that the soluble microenvironment can alter T-cell biomechanics in addition to F-actin levels, immunological synapse formation, and cancer cell killing. We found that when mechanical forces were reduced, T-cells lost most of their effector functions including cytokine and cytolytic mediator production, TCR triggering, immunological synapse formation, and cytotoxicity. The reduced force signal could be a result of a reduced magnitude of TCR forces or a reduced frequency of TCR forces and our results do not distinguish between these two possibilities. Additionally, our prior work showed that TCR forces can be dampened either by using an irreversible surface tether that detaches from the surface upon force exertion and terminates TCR forces, or by using a cytoskeletal inhibitor, which both lead to dampened TCR triggering and T-cell function^25^. Thus, the observed reduction in TCR force and corresponding reduction in T-cell activation is consistent with past work. Conversely, past work has shown that applying exogenous forces to the TCR-antigen complex can enhance TCR triggering, which was observed using AFM^24^, optical tweezer^26^, and BFP^27,30^ measurements. Here we note that SCM leads to enhanced TCR-pMHC force as well as increased T-cell function, which is consistent with prior literature.

The FDA-approved drug blinatumomab harnesses endogenous T-cell function by bridging CD3+ T-cells and CD19+ B-cells to promote immunological synapse formation and cytotoxicity^75^. Using DNA-based tension probes, we discovered that T-cells exert a quantifiable mechanical force to blinatumomab. Additionally, we discovered that the soluble microenvironment modulates mechanical forces exerted by T-cells on blinatumomab. We showed that decreased mechanical forces through blinatumomab are correlated with poor immunological synapse formation and low levels of cytotoxicity, providing a potential biomarker associated with the function of the drug.

An important underlying question that stems from this work is: what are the secreted factors modulating T-cell gene expression, TCR mechanics, and T-cell function? Based on previous work^45^, we identified top cytokines in SCM and ACM (**Table S1**) and pre-treated OT-1 T-cells prior to seeding them on tension probes. However, there were no significant changes in force signal after cytokine treatment (**Extended Data Fig. 10**). This observation suggests that the effects of conditioned media may be due to other soluble factors, including metabolites such as amino acids, lipids and nucleic acids, or a combination of these factors. Our recent work identified thousands of metabolites in ACM^85^, and this guided us to testing the impact of adenosine, guanosine, and inosine treatment, as well as L-Asp and L-Glu on T-cell F-actin levels and cell spreading (**Extended Data Fig. 11),** but none of these metabolites recapitulated the effects of ACM. Given the thousands of potential secreted metabolites that likely contribute to ACM activity, further future studies are needed to uncover the most potent mediators of this complex response. This will be the topic of future work and is outside the scope of our current report.

Our results support using a biomechanical readout, or “biomechanical biomarkers,” as a novel method to perform quality control on cell-based immunotherapies. Current methods rely on cytokine production and receptor upregulation^86,87^, however, biomechanical biomarkers and TCR avidity studies are just beginning to be explored in this context^88–90^. While our technology has limitations including the use of cell-surface interactions instead of cell-cell interactions, it also offers advantages including the ability to test any ligand of interest in a 96-well plate format. Additionally, our technology only reports a correlation between TCR forces and T-cell function and does not determine a causal effect. However, leveraging T cell biomechanics as an additional quality control marker for cell-based immunotherapies may complement current strategies to lead to more successful immunotherapy treatments in the future.

## Methods

### Materials

200 proof ethanol was purchased from Fischer Scientific (# 04-355-223). Streptavidin (# S000-01) was purchased from Rockland. Hydrogen peroxide (# 216763), sulfuric acid (# SX1244-6), (3-Aminopropyl)triethoxysilane (APTES, # 440140), dimethylsulfoxide (DMSO, # 67-68-5), bovine serum albumin (BSA, # 0735078001), Triton X-100 (# T8787), Fillipin III dye (# F4767), and sodium ascorbate (# A4934) were purchased from Millipore Sigma. mPEG (# MF001023-2K) and lipoic acid PEG (LA-PEG, # HE039023-3.4K) were purchased from Biochempeg. Sulfo-NHS-acetate (# 26777), Pierce 16% methanol free formaldehyde (# 28906), Teflon racks (# C14784), CellTrace Violet and CFSE (# C34571, # C34554), anti-LAMP1-AF488 (# 53-1079-42) anti-WAVE1 (# PA5-78273, goat-anti-rabbit secondary AF647 (A-21245), CD3/CD28 mouse dynabeads (# 11456D), and Live/Dead stains (# L34976, # L34955) were purchased from ThermoFischer Scientific. Tannic acid-modified 8.8 nanometer gold nanoparticles (AuNPs) were custom synthesized by Nanocomposix. Cy3B-NHS (# PA63101) was purchased from GE Life Sciences. P2 gel (# 150-4118 was purchased from BIO-RAD. RPMI 1640 (# 10-041-CV), fetal bovine serum (FBS, # 35010CV), penicillin/streptomycin (# 30-002-CI), and 1× DPBS (# 21-031-CV) were purchased from Corning. Murine CD4+ and CD8+ positive selection kits (# 130-117-043, 130-117-044), human CD3+ positive selection kit (# 130-050-101), and murine CD3+ and CD8+ negative selection kits (# 130-095-130, # 130-104-075) were purchased from Miltenyi Biotec. SiR-Actin (# CY-SC001) was purchased from Cytoskeleton, Inc. RBC Lysis buffer (# 420301), anti-CD3ε-biotin (# 100304), anti-CD3ε (# 100302), anti-CD8a-APC (# 100712), anti-CD69 PE (# 104508), anti-CD25-AF647 (# 102020), anti-IFNγ-PE (# 505807), anti-TNFα-FITC (# 506303), anti-Perforin-APC (# 154303), and anti-Granzyme B-Pacific Blue (# 515407) were purchased from BioLegend. Anti-TCR Vα2-PE (# 553289), anti-CD8α-BUV395 (# 563786), Stain Buffer (FBS, # 554656) and BD Cytofix/Cytoperm kit (# 554714) were purchased from BD Biosciences. Anti-LFA2-iFlour568 (# 100210B0) was purchased from AAT Bioquest. Anti-β-actin (# 4967) and anti-GAPDH (# 97166) were purchased from Cell Signaling Technology. Anti-rabbit-680 (# 926-68071) and anti-mouse-800 (# 926-32210) were purchased from LI-COR. No. 2 Glass Coverslips (# 48382-085) were purchased from VWR. RNeasy mini prep kit (# 74106) was purchased from QIAGEN. TCO-NHS (# BP-22417) and Methyltetrazine-PEG4-Azide (# BP-22446) were purchased from BroadPharm. Blinatumomab (# bimab-hcd19cd3) and SIINFEKL-OVA peptide (# vac-sin) were purchased from InVivoGen. Cupric Sulfate, 5-hydrate (CuSO_4_•5H_2_O # 4844) was purchased from Mallinckrodt chemicals. Monomeric and tetrameric versions of H2-K^b^ OVA 257-264 SIINFEKL pMHC were generously provided by the NIH Tetramer Core Facility.

### Oligonucleotides

All DNA oligonucleotides were purchased from and custom synthesized by Integrated DNA Technologies (Coralville, IA). **Table S1** indicates the names, sequences, and chemical modifications of all DNA oligonucleotides used to form tension probes in this manuscript.

### Mice

All animal work was done at the Division of Animal Resources at Emory University following all Institutional Animal Care and Use Committee protocols. C57BL/6 mice were purchased from The Jackson Laboratory and spleens were harvested between the ages of 6-12 weeks. OT-1 mice (C57BL/6-Tg(TcraTcrb)1100Mjb/J) were purchased from The Jackson Laboratory and bred in the Division of Animal Resources at Emory University. OT-1-Nur77-GFP mice were generously provided by Dr. Byron B. Au-Yeung (Department of Immunology at Emory University). Male and female mice were used equally.

### Cell lines

The human B-cell acute lymphoblastic leukemia (B-ALL) cell line, Nalm6, was generously gifted by Dr. Lia Gore (University of Colorado). Nalm6 cells were maintained in RPMI 1640 (Corning # 10-041-CV) with 10 % FBS (Corning, # 35010CV).

### Equipment

The major equipment used in this study includes a Barnstead nanopure water system (ThermoFischer Scientific), a Nanodrop 2000 UV-Vis Spectrophotometer (ThermoFischer Scientific), a high-performance liquid chromatography (HPLC, Agilent) 1100 with an AdvanceBio Oligonucleotide C18 column (Agilent # 653950-702, 4.6 x 150 mm, 2.7 µm), a Nikon Ti-Eclipse inverted fluorescence microscope equipped with a piezo-controlled motorized stage, ANDOR 512 EMCCD camera, perfect focus system, CFI Apo 100x NA 1.49 oil objective, RICM, DAPI, GFP, TRITC, and CY5 cubes, and driven by NIS Elements software (Nikon), a CytoFLEX flow cytometer (Beckman Coulter), a BD LSR II flow cytometer (BD Biosciences), a BD FACSymphony A5 flow cytometer (BD Biosciences), an Amnis Imagestream^X^ MK II imaging flow cytometer (Luminex), a LI-COR Odyssey CLx gel scanner (LI-COR), and a Bioanalyzer (Agilent, Emory Integrated Genomics Core).

### Preparation of conditioned media

Bone marrow stromal cell and adipocyte conditioned medium was generated as previously reported^45^. Briefly, murine OP-9 bone marrow stromal cells were differentiated into adipocytes as previously described^44,45^. OP-9 cells were plated at 10^5^ cells/well in a 6-well plate in DMEM with 10% FBS. On the next day, the media was changed to either fresh DMEM with 10% FBS for preparing bone marrow stromal conditioned media (SCM) or insulin-oleate media (DMEM supplemented with 1.8 mM Oleate bound to BSA, molar ratio 5.5:1 Oleate:BSA) for adipocyte differentiation to generate adipocyte conditioned media (ACM). Media was harvested from OP-9 bone marrow stromal cells or adipocytes after 3 days of culture for use in experiments. Multiple batches of conditioned media were used throughout this manuscript. Our previous work has shown that different baches of CM have similar metabolite composition^85^. UCM was RPMI 1640 (Corning, # 10-041-CV) with 10% FBS and 100 U/mL penicillin, 100 μg/mL streptomycin.

### Gold nanoparticle surface preparation for murine T-cell probes

Gold nanoparticle DNA probe surfaces were prepared as described previously^25,76,77^. Briefly, No. 2 glass coverslips (VWR # 48382-085) were placed in a Teflon rack (Thermo, # C14784) and immersed and sonicated in Nanopure water for 10 minutes followed by sonication in 200 proof ethanol for 10 minutes. After sonication, the coverslips were baked in an 80°C oven to evaporate all remaining ethanol. Once completely dry, the Teflon rack containing coverslips was immersed in 40 mL freshly made piranha solution (3:1 v/v mixture of H_2_SO_4_:H_2_O_2_) for 30 minutes. CAUTION: piranha solution gets extremely hot and is highly explosive if mixed with organics. After 30 minutes, the slides were washed with 40mL Nanopure water 6 times to remove all piranha solution. Slides were then washed 3 times with 200 proof ethanol to remove all water. Slides were then incubated in 3% APTES in 50mL 200 proof ethanol for 1 hour at room temperature. After incubation, slides were washed 3 times with 200 proof ethanol and baked in an 80°C oven for 30 minutes. The amine-modified coverslips were allowed to cool for 5 minutes and placed in petri dishes lined with parafilm. 200 μL of 0.5% w/v lipoic acid-PEG-NHS and 2.5% w/v mPEG-NHS in 0.1M NaHCO_3_ was added to each coverslip was incubated for 1h at room temperature. Coverslips were rinsed with nanopure water and incubated with 1 mg/mL sulfo-NHS-acetate in 0.1M NaHCO_3_ for 30 minutes to passivate any unreacted amine groups. Coverslips were intensively washed with Nanopure water and then incubated with 500 uL of 20nM gold nanoparticles (8.8 nm, tannic acid modified, custom synthesized by Nanocomposix) for 30 minutes at room temperature. Meanwhile, DNA hairpin probes were assembled in 1M NaCl by mixing the 4.7pN hairpin strand (0.3 μM), Cy3b-ligand strand (0.3 μM), and BHQ2-anchor strand (0.3 μM) at a ratio of 1: 1.1: 1.1. The mixture was annealed by heating to 95°C for 5 minutes and cooling at 25°C for 30 minutes. After annealing, an additional 2.7 μM of the BHQ2 anchor strand was added to the DNA solution for additional quenching. After the 30 minute incubation with gold nanoparticles, coverslips are rinsed extensively with nanopure water, followed by rinsing with 1 M NaCl. Next, the annealed DNA probe and quencher solution was added to the coverslips and incubated overnight in the dark at 4°C. The next day, coverslips were washed with PBS and 40 μg/mL of streptavidin in PBS was added to the functionalized surface for 1h at room temperature. The coverslips were washed with PBS, then 40 μg/mL of biotinylated αCD3ε or OVA-pMHC was added to the functionalized surface for 1h at room temperature. Finally, the slides were washed with PBS, assembled in imaging chambers, and immediately used for imaging.

### TCO surface preparation for blinatumomab tension probes

TCO-glass surfaces were prepared as described previously^91^. Coverslips were prepared identical to the AuNP coverslips through the APTES step.100 μL of 4 mg/mL TCO-NHS in DMSO was added to each amine-functionalized coverslip and another coverslip was placed on top to create a “sandwich.” Coverslips were incubated for at least 12h or up to 1 week at RT.

Coverslips were intensively washed with ethanol then PBS to remove all DMSO, and then incubated with 0.1% BSA in PBS for 30 min to block any nonspecific interactions. Meanwhile, DNA hairpin probes were assembled in 1M NaCl by mixing the 4.7pN hairpin strand (0.2 μM), Cy3b-ligand strand (0.22 μM), and BHQ2-anchor strand (0.22 μM) at a ratio of 1: 1.1: 1.1. The mixture was annealed by heating to 95°C for 5 minutes and cooling at 25°C for 30 minutes. After the 30 minute incubation with 0.1% BSA, coverslips are rinsed extensively with PBS. Next, the annealed DNA probe solution was added to the coverslips and incubated for 1h at RT. Next, coverslips were washed with PBS and 40 μg/mL of streptavidin in PBS was added to the functionalized surface for 1h at room temperature. The coverslips were washed with PBS, then 10 μg/mL of biotinylated CD19 protein (Acro Biosystems, # CD9-H82E9) was added to the functionalized surface for 1h at room temperature. Next, 10 ug/mL blinatumomab (InVivoGen, # bimab-hcd19cd3) in PBS was added to CD19 probes for 1h. Finally, the slides were washed with PBS, assembled in imaging chambers, and immediately used for imaging.

### Labeling DNA ligand strand with Cy3B via NHS coupling

Oligonucleotides were custom synthesized by IDT to include a terminal amine for NHS-Cy3b coupling. Briefly, an excess of Cy3b-NHS dye (50 μg, GE Life Sciences, # PA63101) was dissolved in DMSO and added to a solution containing 10nmol of the DNA ligand strand in 1xPBS with 0.1M NaHCO_3_. The reaction incubated at room temperature overnight. The next day, byproducts, salts, and unreacted Cy3B-NHS was removed via P2 gel (BIO-RAD, #150-4118) filtration in a Nanosep MF centrifugal device. The DNA-Cy3B conjugate was further purified by reverse-phase HPLC with an Agilent AdvanceBio Oligonucleotide C18 column (653950-702, 4.6 x 150 mm, 2.7 μm).

### Labeling DNA anchor strand with tetrazine using a copper click reaction

Tetrazine labeling of oligonucleotodies was done as described previously^91^. Oligonucleotides were custom synthesized by IDT to include a BHQ2 quencher and a terminal alkyne for azide-tetrazine coupling. Briefly, alkyne-modified DNA was reacted with an excess amount of Methyltetrazine-PEG4-Azide (Broadpharm, # BP-22446) in the presence of CuSO_4_ (0.20 μmol), sodium ascorbate (0.50 μmol), and THPTA (0.25 μmol) in 40 μL (1:1 ratio of DMSO: 18.2 MΩ MilliQ water) for 1 hour at 50°C. The product was then filtered using P2 gel filtration in a Nanosep MP centrifugal device. The tetrazine-DNA conjugate was further purified by reverse-phase HPLC with an Agilent AdvanceBio Oligonucleotide C18 column.

### CD4+ and CD8+ polyclonal murine T-cells on tension probes

Polyclonal CD4+ and CD8+ T-cells were isolated from C57BL/6 mice, which are housed and bred at the Division of Animal Resources at Emory University following all Institutional Animal Care and Use Committee protocols. CD4+ and CD8+ T-cells were isolated from spleens using CD4+ or CD8+ positive selection according to the manufacturer’s protocol (Miltenyi Biotec, # 130-117-043, 130-117-044). T-cells were resuspended in UCM, SCM, or ACM at 1×10^6^ cells/mL in a 24 well plate and incubated at 37°C, 5% CO_2_ for 24 h prior to seeding on tension probes. Images were taken 30 minutes after adding cells to tension probes. Fluorescence intensity values of individual cells were quantified from 20 images in FIJI (ImageJ) software.

### CD3+ T-cells from a murine obesity model on tension probes

CD3+ T-cells were isolated from control or DIO C57BL/6 mice, which were generated using methods previously described^45^. CD3+ T-cells were isolated using negative selection according to the manufacturer’s protocol (Miltenyi Biotec, # 130-095-130). T-cells were then resuspended in RPMI + 10% FBS and 200,000 cells were seeded per tension probe surface. Images were taken 30 minutes after adding cells to tension probes. Fluorescence intensity values of individual cells were quantified from 20 images in FIJI (ImageJ) software.

### RNA sequencing of primary CD3+ murine T-cells

Polyclonal CD3+ T-cells were isolated from C57BL/6 mice, which are housed and bred at the Division of Animal Resources at Emory University following all Institutional Animal Care and Use Committee. CD3+ T-cells were isolated from spleens using CD3+ negative selection according to the manufacturer’s protocol. (Miltenyi Biotec, # 130-095-130). CD3+ T-cells were resuspended in UCM, SCM, or ACM at 1×10^6^ cells/mL in a 24 well plate and incubated at 37°C, 5% CO_2_ for 4 h prior to RNA isolation. RNA was extracted using the QIAGEN RNeasy Mini Kit following the manufacturer’s protocol (QIAGEN, # 74106) and quality control was performed using a Bioanalyzer (Agilent).

Isolated RNA samples were submitted to the Emory Integrated Genomics Core (EIGC) for library preparation and sequencing. cDNA was generated using SMART-seq v4 Ultra Low Input RNA Kit (Takara Bio) and the final sequencing library was made using the Nextera XT Kit (Illumina) following standard protocols. cDNA libraries were sequenced on a Novaseq S1 flowcell that provides >50M reads per sample. Raw reads were accessed through the EIGC, in Unix, and quality checked using FastQC. Adaptor sequences and low-quality reads were removed with Trimmomatic. Reads were aligned to GRCm39 in Bioconductor (in R) using a ‘seed-and-vote’ algorithm (Rsubread1 package). Count matrices were generated with “feature Counts” in Rsubread. DESeq2 (in R) was used to generate normalized RNA-seq signal values (median of ratios method to account for sequencing depth), to identify differentially expressed genes (log2 fold change ≤ -2 or ≥ 2, adjusted p value ≤ 0.05) and to generate a PCA plot (single value decomposition). Agglomerative clustering was performed using the hclust function. Distances between observations were measured in Euclidean distances using complete linkage clustering. Prism 9 (GraphPad) software was used to generate heatmaps and bar charts of normalized RNA-seq signal values. For the heatmaps, percent maximum expression was calculated in Prism9 with highest value in each row set as 100%.

### OT-1 murine T-cells on tension probes

OT-1 T-cells are primary mouse CD8+ T cells that specifically recognize the ovalbumin peptide residues 257-264 (SIINFEKL) presented on a class I MHC. OT-1 transgenic mice were housed and bred at the Division of Animal Resources at Emory University following all Institutional Animal Care and Use Committee protocols. OT-1 T-cells were magnetically isolated from spleens using CD8a+ negative selection according to the manufacturer’s protocol (Miltenyi Biotec, # 130-104-075). Purified OT-1 T-cells were resuspended in UCM, SCM, or ACM at 1×10^6^ cells/mL in a 24 well plate and incubated at 37°C, 5% CO_2_ for 24h prior to being seeded on tension probes. Images were taken 15 minutes after seeding OT-1 T-cells on tension probes. Fluorescence intensity values of individual cells were quantified from at least 10 images in FIJI (ImageJ).

### SirActin, WAVE1, and Filipin III staining of T-cells

OT-1 T-cells were isolated as previously described and resuspended in UCM, SCM, or ACM at a concentration of 1×10^6^ cells/mL in a 24 well plate and incubated at 37°C, 5% CO_2_ for 24h prior to being seeded on antiCD3ε-coated glass bottom 96 well plates. T-cells from control or DIO mice were isolated as described above. Glass bottom plates were prepared by incubating with 10μg/mL aCD3ε in PBS at 37°C for 1h, then blocking with 0.5% BSA in PBS at 37°C for 1h. Wells were washed with PBS prior to cell addition. Cells treated with UCM, SCM, or ACM, or cells from control or DIO mice were added to wells and incubated at 37°C, 5% CO_2_ for 30 minutes, and immediately fixed with 4% methanol-free formaldehyde (Thermo, # 28906) in PBS. For Filipin III staining, cells were stained with 50 μg/mL Fillipin III dye (Millipore Sigma, # F4767) for 30 min at 4°C and 3 times with 1xPBS prior to imaging. For SirActin and WAVE1 staining, cells were permeabilized after fixation with 0.1% Triton-X100 (Millipore Sigma, # T8787) in PBS for 3 minutes, washed with PBS, then blocked with 2% BSA in PBS for 1h at room temperature. For SirActin, wells were washed with PBS, then 0.5 μM SirActin (Cytoskeleton, Inc, # CY-SC001) in 1% BSA in PBS was added to each well for 30 minutes. Cells were imaged without washing the SirActin dye out of the wells following the manufacturer’s protocol. For WAVE1 staining, wells were washed with with PBS and αWAVE1 (ThermoFisher Scientific, # PA5-78273) was added at a 1:100 dilution in PBS containing 0.5% BSA and 0.01% Triton-X 100 for 1 hr at RT. Wells were washed 3x with PBS for 5 min each. Next, a secondary fluorescent antibody (ThermoFisher Scientific, # A-21245) was added at a concentration of 4 μg/mL in PBS containing 0.5% BSA and 0.01% Triton-X 100 for 1 hr at RT. Cells were washed 3x for 5 min each with PBS. Cells were immediately imaged. Fluorescence intensity values of individual cells were quantified from at least 10 images in FIJI (ImageJ) software.

### Western blotting

OT-1 T-cells were isolated as previously described^45,92^ and resuspended in UCM, SCM, or ACM at a concentration of 1×10^6^ cells/mL in a 24 well plate and incubated at 37°C, 5% CO_2_ for 24h prior to being lysed. 6×10^6^ cells per condition were lysed and prepared following a previously established method^92^. Lysates were probed using anti-β-actin and anti-GAPDH (1:1000 dilution, Cell Signaling Technology, # 4967, # 97166). Either anti-rabbit-680 or anti-mouse-800 (1:5000 dilution, LI-COR, # 926-68071, # 926-32210) were used as the secondary antibodies. Protein signals were detected using a LI-COR Odyssey CLx and protein quantification via signal intensity was performed using FIJI (ImageJ) software.

### Flow cytometry on unstimulated OT-1 T-cells

OT-1 T-cells were isolated as previously described and resuspended in UCM, SCM, or ACM at a concentration of 1×10^6^ cells/well in a 24 well plate and incubated for 24h at 37°C, 5% CO_2_. For extracellular staining, cells were washed once in Stain Buffer (BD Biosciences, # 554656) and stained with either anti-CD8α-APC or anti-TCR Vα2-PE (BioLegend # 100712, BD Biosciences, # 553289) for 45 min, then washed two times with Stain Buffer. Cells were then fixed in fixation solution (BD Biosciences, # 554714), washed two times, and resuspended in Stain Buffer and kept on ice until running flow cytometry. Flow cytometry was performed on a Beckman Coulter CytoFLEX and the data was analyzed using FlowJo.

### OT-1 72 h activation and flow cytometry

Ovalbumin (OVA)-specific CD8+ T-cells were purified from OT-1 C57BL/6 mice using magnetic-activated cell sorting (MACS). OT-1 T-cells (5 × 10^4^ cells/well) were co-cultured with 0.5 ng/mL SIINFEKL peptide (InVivoGen, # vac-sin)-pulsed splenocytes (5 × 10^4^ cells/well) in 96-well flat bottom plates containing RPMI 1640 media supplemented with 10% FBS. On day 3 of culture, cells were harvested, and tetramer staining was performed to identify OVA-specific CD8+ T-cells. The production of cytokines and cytolytic mediators was determined in OT-1 T-cells using intracellular cytokine staining as previously described^93,94^ to determine interferon-gamma (anti-IFN-γ PE, Biolegend, # 505807), tumor necrosis factor-alpha (anti-TNF-α FITC, Biolegend, # 506303), perforin (anti-Perforin APC, Biolegend, # 154303), and granzyme B (anti-Granzyme B Pacific Blue, Biolegend, # 515407) production using flow cytometry (% positive T-cells). All samples were run on the CytoFLEX (Beckman Coulter) flow cytometer and data were analyzed using the FlowJo Version 9 software.

### OT-1 2 h activation and flow cytometry

Nur77-GFP-expressing OT-1 T-cells were treated for 20 h in UCM or 20% SCM or ACM at 37°C, 5% CO_2._ T-cells were then co-cultured with splenic antigen presenting cells at a 2:1 ratio of APCs:T-cells in the presence of 0.1 μM SIINFEKL peptide. After stimulation, T-cells were stained with Live/Dead stain (Thermo, # L34796 or L34995), anti-CD69-PE (Biolegend, # 104508), anti-CD25-AF647 (BioLegend, # 102020), and anti-CD8α-BUV395 (BD Biosciences, # 563786) prior to running flow cytometry. T-cell populations were assessed for the percent of live CD8+ cells that were Nur77^HI^CD25^HI^ or CD69^HI^CD25^HI^. All samples were run on either a BD LSR II or a BD FACSymphony A5 (BD Biosciences) and data was analyzed using FlowJo.

### *Ex vivo* T cell stimulation from lean and obese mice

Lean and obese mice were generated as previously described using the diet-induced model of obesity^45^. After 3 months of feeding control or high-fat diets, splenic-derived CD4^+^ and CD8^+^ T-cells were isolated via FACs sorting. Purified T-cells (5 x 10^4^ cells/well) were stimulated *in vitro* using αCD3/αCD28 stimulation beads (ThermoFisher Scientific, # 11456D) at 1 µg/mL and 5 µg/mL, respectively. After 3 days of culture, stimulation beads were removed using the DynaMag™-2 magnet. Intracellular staining was performed on purified T-cells as previously described^93–95^ to ascertain the production of IFN-γ, TNF-α, perforin, and granzyme B. All flow samples were acquired on the Cytek Aurora 5 (BD Biosciences and analyzed using FlowJo v. 10.0.1 software (Ashland, Oregon).

### Human PBMC isolation

PBMCs were isolated from de-identified, normal donor buffy coats purchased from ZenBio using a ficoll-density gradient. CD3+ T-cells were positively selected from PBMCs via magnetic separation (Miltenyi Biotech, CD3 Microbeads, # 130-050-101). T-cell purity was assessed at > 90% CD3+ of Live cells, and the cells were cryopreserved. Cells were cultured in RPMI (10% FBS, 100 U/mL penicillin, 100 μg/mL streptomycin) at 37°C, 5% CO2 unless otherwise specified.

### Human T-cells on blinatumomab tension probes

CD3+ T-cells were thawed and incubated overnight at 37°C, 5% CO_2_ to recover. Next, CD3+ T-cells were cultured in UCM, SCM, or ACM at a density of 5×10^5^ cells/mL for 24 h at 37°C, 5% CO_2_ prior to seeding on blinatumomab tension probes. CD3+ T-cells were allowed to interact with the tension probes for 15 minutes prior to adding the locking strand. The locking strand was added at a concentration of 1 μM for 10 minutes prior to imaging. In brief, the locking strand is essential to boost signal when the tension signal is weak or transient. This helps with visualizing the signal and also in quantifying the number of mechanical events of a specific magnitude^76,77^. We find that cryopreserved T-cells isolated from PBMCs tend to display weaker tension signal compared T-cells freshly isolated from mouse splenocytes, therefore why we used the locking strand. Fluorescence intensity values of individual cells were quantified from 15 images in FIJI (ImageJ) software.

### T-cell immunological synapse formation with blinatumomab

Nalm-6 B-ALL cells (stained with CellTrace Violet, Thermo # C34571) and CD8+ T-cells were treated separately for 24 h in UCM, SCM, or ACM. After initial treatment, cells were replated at a 1:1 ratio of Nalm-6:T-cell and co-cultured in the presence of blinatumomab (7 ng/mL) or vehicle (1x PBS). After a 24 h co-culture, cells were directly fixed in culture with formaldehyde, permeablized with BD perm/wash, and stained with anti-LAMP-1 AF488 (Thermo # 53-1079-42), anti-LFA-2 iFlour568 (AAT Bioquest, # 100210B0), and SiR-Actin (Cytoskeleton, Inc # CY-SC001). Samples were run on Amnis Imagestream^X^ MK II at 60x in EDF mode and data was analyzed in IDEAS software.

In IDEAS software, potential conjugates were distinguished from single cells using the area vs. aspect ratio of brightfield images. Conjugates double positive for CellTrace Violet (Nalm6) and LFA2 AF568 (T-cell) were gated and further refined by conjugates containing only one CTV+ cell and one LFA2+ cell. A gate limiting centroid distance between a CTV+ Nalm6 and an LFA2+ T-cell was then incorporated to eliminate images of CTV+LFA2+ conjugates that were captured simultaneously but either spatially separate or overlapping rather than interfacing via brightfield. CTV+LFA2+ conjugates in contact were then identified as synapsing by defining a mask covering the interface between the Nalm6 and T-cell and gating on concentrated positive actin signal (max pixel feature) within the contact surface between the Nalm6 and T cell.

### T-cell cytotoxicity assay with blinatumomab

Nalm-6 and CD3+ T-cells were stained with tracking dyes CellTrace Violet (Thermo, # C34571) and CellTrace CFSE (Thermo, # C34554), respectively, prior to culture. T-cells were independently pre-treated with UCM, SCM, or ACM for 24h. Both T-cells and Nalm6 cells were counted and then plated in a 96-well u-bottom plate at a 1:1 E:T ratio in RPMI, SCM, or ACM and treated with vehicle (PBS) or 3.5ng/mL blinatumomab (InVivoGen, # bimab-hcd19cd3) for 24h and 48h. The cells were then prepared for flow cytometry. Primary cells were stained for purity post-isolation using anti-CD3 (OKT3, BioLegend, # 317301), and Near IR Live/dead stain (Invitrogen, Thermo, # L3495). Following 24h and 48h co-culture, cells were stained with Near IR Live/Dead. All samples were run on a Cytoflex machine (Emory Pediatric Flow Cytometry Core) and analyzed with FlowJo 2 software. Viability measured by flow cytometry was used to calculate % lysis with the following equation: % Lysis = 100 x (%Treated Sample [Dead Nalm6] - %Control Sample [Dead Nalm6]).

### Image acquisition

All images were taken on a fully motorized Nikon Ti-E Eclipse inverted fluorescence microscope driven by NIS Elements software. The microscope features used in these experiments were a CFI Apo 100x NA 1.49 oil objective (Nikon), an ANDOR 512 EMCCD camera, a piezo-controlled stage, RICM, DAPI, TRITC, and CY5 cubes, and a perfect focus system to allow the capture of multipoint images without losing focus. Imaging was performed at room temperature in UCM, SCM, or ACM. Images were taken within 30 minutes after adding T-cells to the surface.

### Image analysis

All image analysis was performed in Fiji is Just ImageJ (FIJI) software. To measure the fluorescence intensity of single cells, a region of interest (ROI) was drawn around the outline of each cell using the RICM channel, and double-checked to make sure it lined up with the fluorescence channel. Next, the local background near the cell was subtracted from the fluorescence channel, and the fluorescence intensity of the previously drawn ROI was measured and recorded. (**Extended Data Fig. 12**) This method was used for tension probe signal, SirActin staining, and cholesterol staining.

### Statistics

Student’s T-tests were performed when comparing two groups and ordinary one-way ANOVAs with post-hoc multiple comparisons were performed when comparing three groups with α = 0.05. Statistical analyses were performed in GraphPad Prism Version 9.

## Acknowledgements

Funding: A.V.K. and R.H. acknowledge Ruth L. Kirschstein Predoctoral Individual National Research Service Awards (NIH NCI F31CA243502 and NIH NCI F31CA243472). J.A.G.H. acknowledges The American Society of Hematology (ASH) Minority Hematology Graduate Award (Grant No. 00092385). R.L.B. acknowledges NIH grant no. R01GM131099-01S1. E.C.D. acknowledges the St. Baldrick’s Foundation (Research Grant 641261) and Emory School of Medicine (Imagine Innovate and Impact (I3), a gift from Woodruff Fund Inc. and through the Georgia CTSA NIH award UL1-TR002378). C.J.H. acknowledges the Mark Foundation for Cancer Research (Grant No. 18-031-ASP), American Cancer Society (Grant No. 00074211), the Transdisciplinary Research on Energetics and Cancer (TREC; Grant No. R25CA203650), Swim Across America (Grant No. 01007), and the Emory Center for Pediatric Cellular Therapies (Grant No. 00103927). K.S. acknowledges NIH grant no. R01GM131099 and R01GM124472.

All schematics throughout this manuscript were created in BioRender.com through a fully licensed subscription (user: Anna V. Kellner). Research reported in this publication was supported in part by the Emory University Integrated Genomics Core (EIGC) of the Winship Cancer Institute of Emory University and NIH/NCI under award number, 2P30CA138292-04. Additionally, research reported in this publication was supported in part by the Pediatrics/Winship Flow Cytometry Core of Winship Cancer Institute of Emory University, Children’s Healthcare of Atlanta and NIH/NCI under award number P30CA138292. The content is solely the responsibility of the authors and does not necessarily represent the official views of the National Institutes of Health. We thank the NIH Tetramer Core Facility (contract number 75N93020D00005) for providing OVA-specific monomers and tetramers. Lastly, we would like to acknowledge Dr. Sunil S. Raikar for helpful discussions.

## Author contributions

A.V.K., C.J.H, and K.S. designed the study and wrote the manuscript. A.V.K., R.H., P.D., J.E., D.K.G., and R.S.A. performed experiments and analyzed data. M.L. J.A.G.H., R.L.B., and R.M. assisted with experiments and experimental design. M.L, J.A.G.H. and A.J.R. provided cells and reagents. M.J. and K.A.H. performed RNA-sequencing analysis. C.C.P, E.C.D, B.B.A-Y, and K.A.H. aided in experimental design and provided guidance for the study. C.J.H. and K.S. contributed to experimental interpretation and provided financial support for the study.

