## Extended Data Figures and Tables for "The T-cell niche tunes immune function through modulation of the cytoskeleton and TCR-antigen forces"

**Table S1 | Previously reported cytokine and chemokine concentrations in SCM or ACM (Lee et al. 2022**)[44]. Values were obtained by Luminex analysis using the Miliplex 30-plex cytokine/chemokine assay kit (Millipore Sigma, # MCYTMAG-70K-PX32). Values represent the concentration in pg/mL ± standard error.

| Cytokine | SCM concentration (pg/mL) | ACM concentration (pg/mL) |
|---|---|---|
| IL-13 | 0 | 2.5 ± 7 |
| IL-15 | 0 | 2.0 ± 5 |
| IP-10 | 32 ± 11 | 39 ± 4 |
| IL-5 | 14 ± 2 | 3 ± 4 |
| VEGF | 123 ± 17 | 55 ± 8 |
| LIF | 19 ± 6 | 2 ± 1 |
| IL-6 | 25 ± 18 | 98 ± 32 |
| TNF-α | 17 ± 5 | 71 ± 12 |
| **Chemokine** | **SCM Concentration (pg/mL)** | **ACM Concentration (pg/mL)** |
| KC | 0 | 3700 ± 1567 |
| LIX | 0 | 130 ± 125 |
| MIP-1α | 0 | 6 ± 5 |
| MIP-2 | 21 ± 2 | 38 ± 37 |
| IP-10 | 30 ± 9 | 36 ± 8 |
| RANTES | 50 ± 16 | 60 ± 22 |

**Table S2 | DNA oligonucleotide sequences for creating DNA-based tension probes.** All DNA oligonucleotides were custom synthesized by Integrated DNA Technologies (Coralville, IA). Table S2 indicates the names, sequences, and chemical modifications of all DNA oligonucleotides used to form tension probes in this manuscript.

| **Sequence Name** | **Description** | **Sequence (5' to 3')** |
|---|---|---|
| 4.7 pN hairpin | Hairpin that has an $F_{1/2}$ = 4.7 pN | GTG AAA TAC CGC ACA GAT GCG TTT GTA TAA ATG TTT TTT TCA TTT ATA CTTTAA GAG CGC CAC GTA GCC CAG C |
| 12 pN hairpin | Hairpin that has an $F_{1/2}$ = 12 pN | GTG AAA TAC CGC ACA GAT GCG TTT GGG TTA ACA TCT AGA TTC TATTTT TAG AAT CTA GAT GTT AAC CCT TTA AGA GCG CCA CGT AGC CCA GC |
| 19 pN hairpin | Hairpin that has an $F_{1/2}$ = 19 pN | GTG AAA TAC CGC ACA GAT GCG TTT CGC CGC GGG CCG GCG CGC GGT TTT CCG CGC GCC GGC CCG CGG CGT TTA AGA GCG CCA CGT AGC CCA GC |
| Ligand strand | Amine-modified top strand that presents the ligand | /5AmMC6/ - CGC ATC TGT GCG GTA TTT CAC TTT - /3Bio/ |
| Cy3b-ligand strand | Ligand strand after in-house fluorophore conjugation | Cy3B - CGC ATC TGT GCG GTA TTT CAC TTT - /3Bio/ |
| BHQ2-anchor strand for AuNP surfaces | Bottom strand that anchors to AuNPs and has a quencher | /5-ThioC6-5/ - TTT GCT GGG CTA CGT GGC GCT CTT - /3BHQ_2/ |
| BHQ2-anchor strand for TCO-MeTz surfaces | Bottom strand that will be functionalized with a methyltetrazine-azide and has a quencher | /5Hexynyl/ - TTT GCT GGG CTA CGT GGC GCT CTT - /3BHQ_2/ |
| MeTz-functionalized BHQ2-anchor strand for TCO-MeTz surfaces | Bottom strand after in-house methyltetrazine conjugation | /5MeTz/ - TTT GCT GGG CTA CGT GGC GCT CTT - /3BHQ_2/ |
| Locking strand | Complementary strand that locks the 4.7 pN hairpin open | AAA AAA CAT TTA TAC |

### Supplementary movie captions

**Supplementary movie 1: Timelapse videos of OT-1 T-cells treated with UCM on 4.7 pN tension probes.** T-cells were pre-treated with UCM for 24 hr, then were seeded on 4.7 pN tension probes presenting the OVA-pMHC for 5 min before timelapse acquisition. Scale bar = 10 μm.

**Supplementary movie 2: Timelapse videos of OT-1 T-cells treated with SCM on 4.7 pN tension probes.** T-cells were pre-treated with SCM for 24 hr, then were seeded on 4.7 pN tension probes presenting the OVA-pMHC for 5 min before timelapse acquisition. Scale bar = 10 μm.

**Supplementary movie 3: Timelapse videos of OT-1 T-cells treated with ACM on 4.7 pN tension probes.** T-cells were pre-treated with ACM for 24 hr, then were seeded on 4.7 pN tension probes presenting the OVA-pMHC for 5 min before timelapse acquisition. Scale bar = 10 μm.

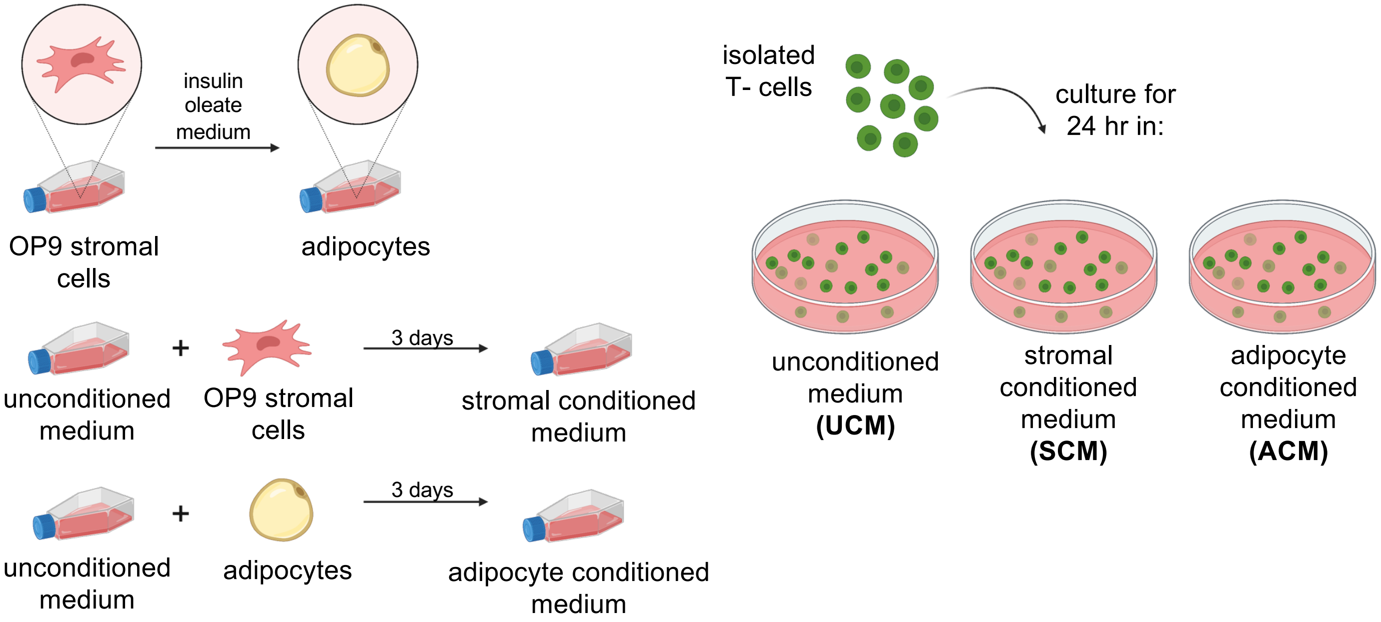


**Extended Data Fig. 1 | Schematic illustration showing the preparation of conditioned media.** OP9 bone marrow stromal cells are differentiated into adipocytes using insulin oleate medium. SCM and ACM are made by incubating unconditioned medium on OP9 stromal cells or adipocytes for 3 days, respectively. T-cells are pre-incubated in each medium prior to every experiment.

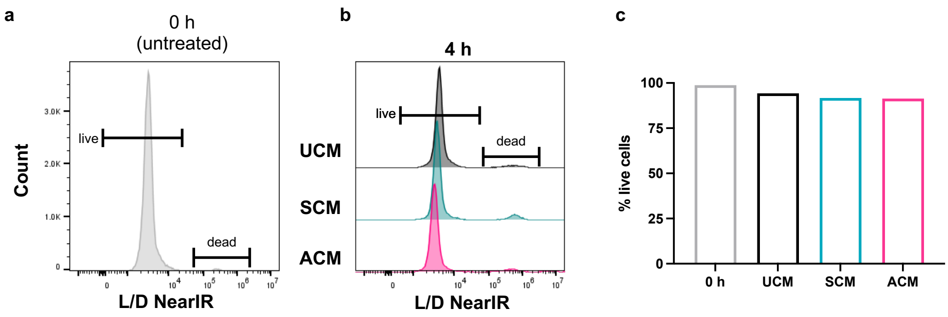


**Extended Data Fig 2 | T cells have high viability after 4 h of treatment with UCM, SCM, or ACM. a**, Viability of freshly isolated, untreated T-cells and **b**, T-cells treated with UCM, SCM, or ACM for 4 h using a NearIR live/dead (L/D stain). **c**, Quantification of T cell viability determined by gating on the live cells.

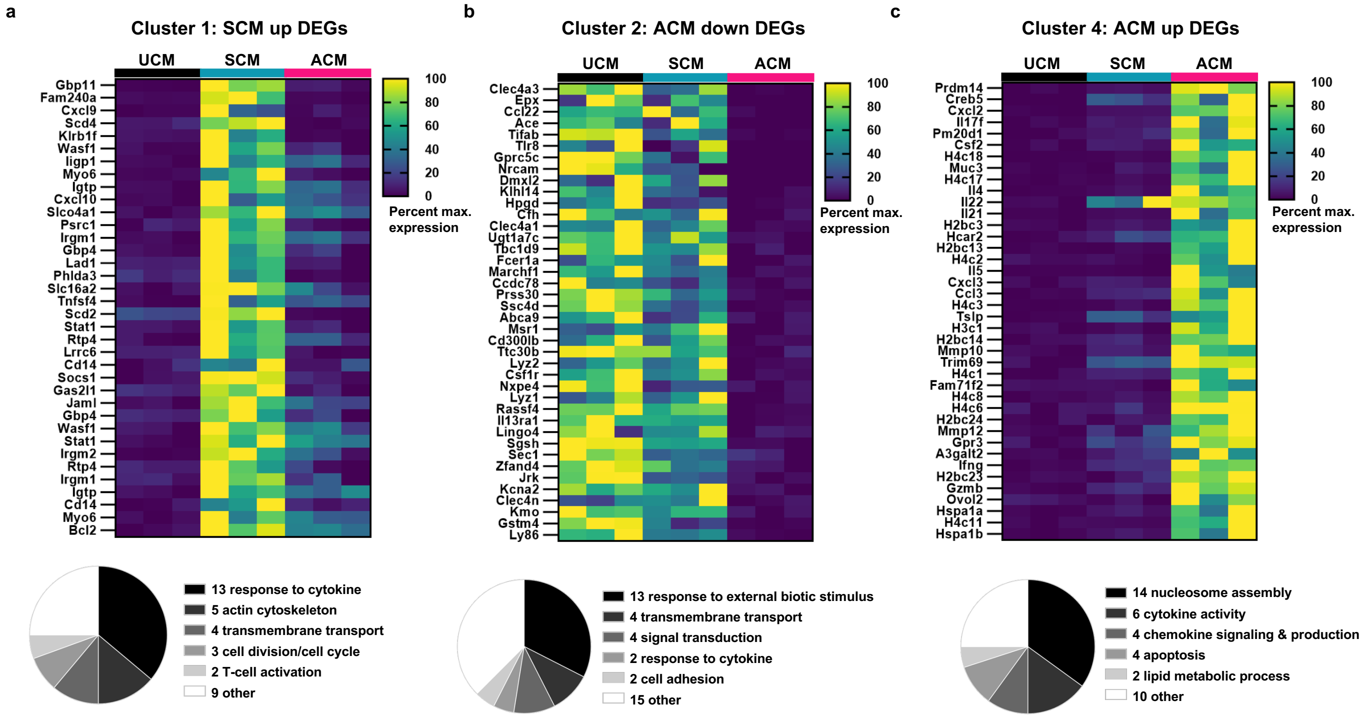


**Extended Data Fig. 3 | Biological processes and molecular functions associated with T-cell DEGs in hierarchical cluster Groups 1, 2, and 4.** Heat maps show row-normalized gene expression values from RNA-seq analysis. Pie charts show the distribution of manually searched biological processes and molecular functions within each group. **a,** Cluster 1 contains 36 DEGs, which are upregulated in SCM versus UCM (log2 FC = 1.36 to 4.86, padj. < 0.05). **b,** The top 40 DEGs in Cluster 2 which are downregulated in ACM versus UCM (log2 FC = -7.9 to -3.3, padj. < 0.05). **c,** The top 40 DEGs in Cluster 4 which are upregulated in ACM versus UCM (log2 FC = 3.73 to 6.56, padj. < 0.05).

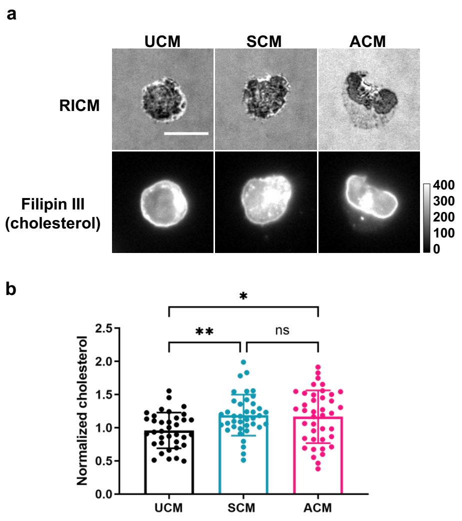


**Extended Data Fig. 4 | The soluble microenvironment alters T-cell cholesterol levels.** **a**, Representative images and **b**, quantification of cholesterol levels OT-1 T-cells treated with UCM, SCM, or ACM for 24 hr. OT-1 T-cells were stained with Fillipin III dye to measure total cholesterol levels. Scale bar = 10 μm. Data points represent fluorescent intensity values of individual cells. Values were normalized to the mean of UCM in each replicate. Graphs represent means ± SD. Statistics were calculated using a one-way ANOVA with a Tukey's multiple comparisons test. UCM v. SCM $^{**}P$ = 0.0093, UCM v. ACM $^{*}P$ = 0.0212, SCM v. ACM $P$ = 0.9465. $n$ = 2 independent animals.

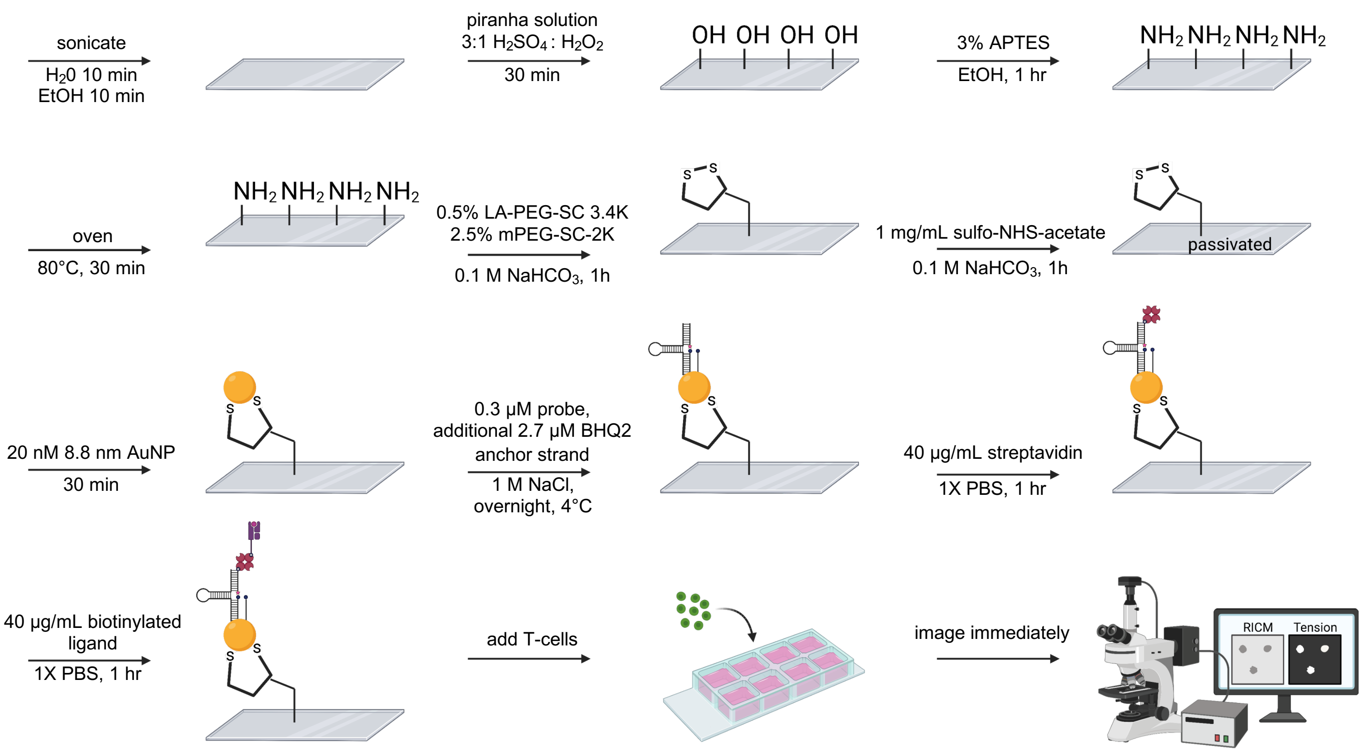


**Extended Data Fig. 5 | Workflow for preparing gold nanoparticle (AuNP)-based DNA tension probes.** All murine T-cell experiments used tension probes that were prepared following this method.

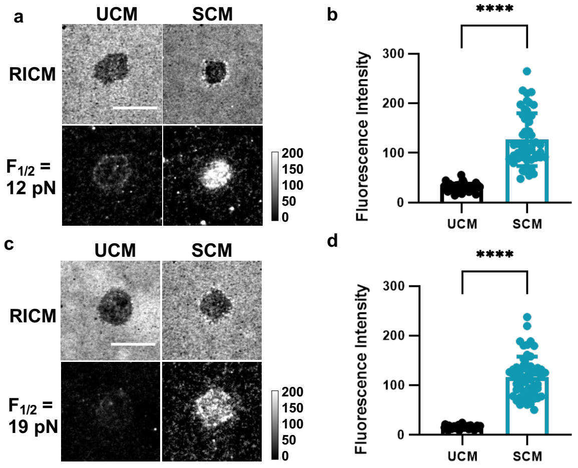


**Extended Data Fig. 6 | SCM increases the magnitude of TCR tension. a**, Representative images and **b**, quantification of 12 pN tension exerted by OT-1 T-cells on OVA-pMHC after 24 hr treatment with UCM or SCM. **c**, Representative images and **d**, quantification of 19 pN tension exerted by OT-1 T-cells on OVA-pMHC after 24 hr treatment with UCM or SCM. Scale bars = 10 µm. Graphs represent means ± SD and data points represent fluorescence intensity measurements of individual cells. Statistics were calculated using a two-tailed student's t-test. ****P < 0.0001, *n* = 3 independent animals.

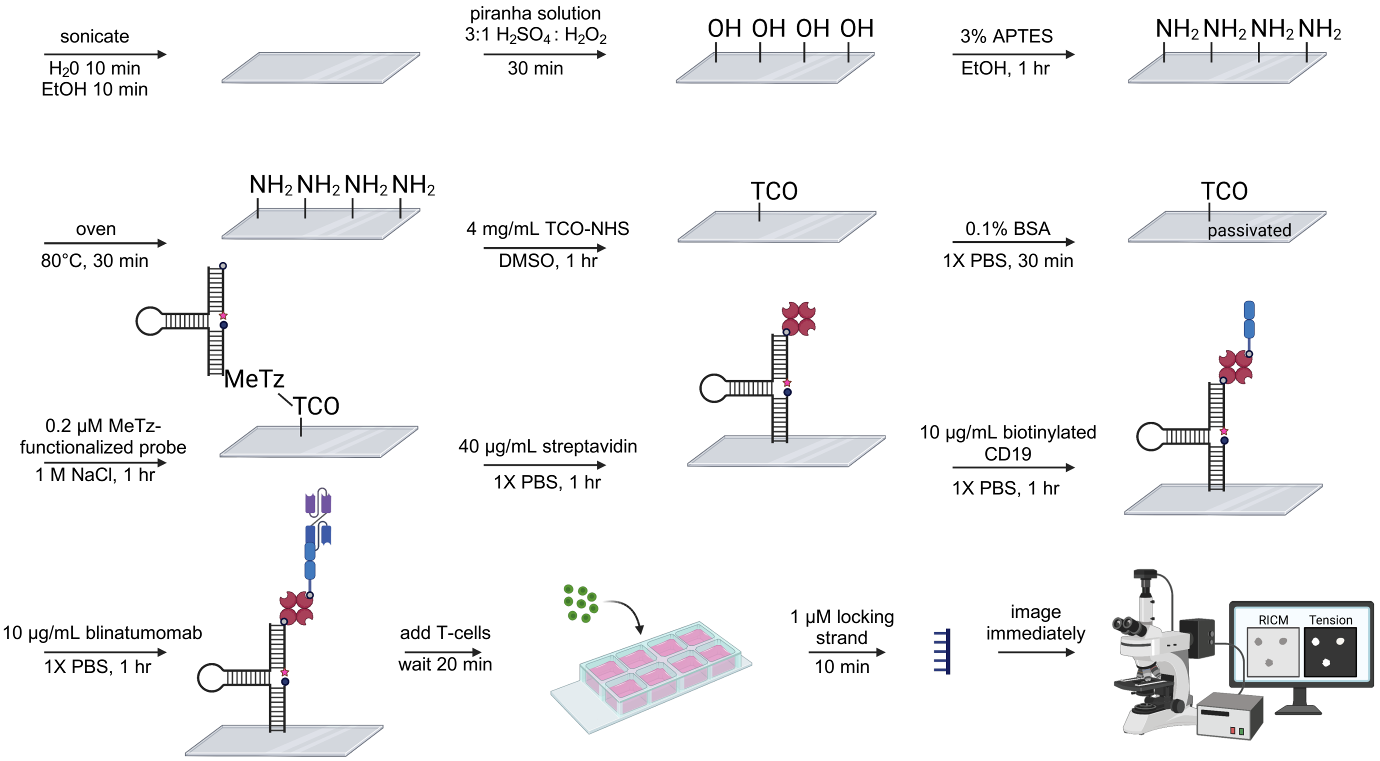


**Extended Data Fig. 7 | Workflow for preparing methyltetrazine (MeTz)-trans-cyclooctene (TCO)-based DNA tension probes**. This method was used for all human donor T-cell experiments.

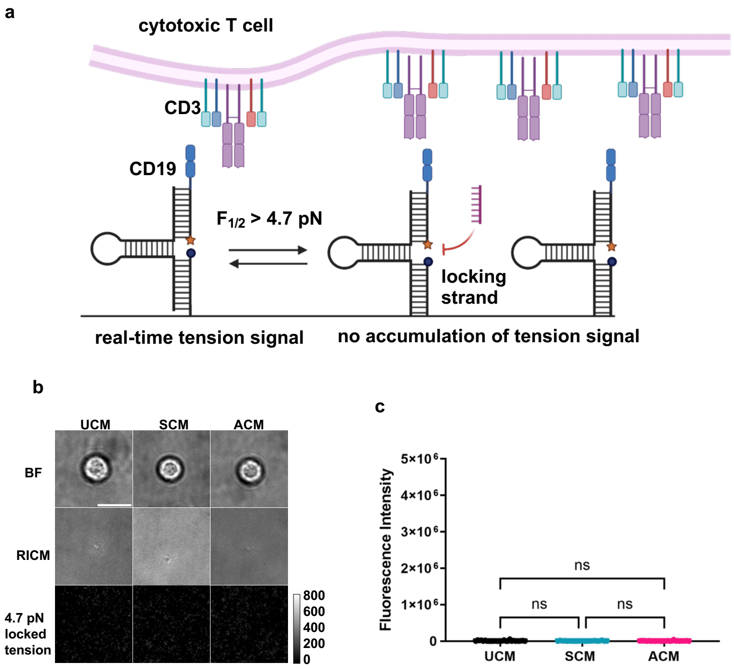


**Extended Data Fig. 8 | Human T-cell binding and tension are specific to blinatumomab. a**, Schematic illustration of a human T-cell failing to interact with and exert a force on a CD19-only presenting tension probe. **b**, Representative images and **c**, quantification of 4.7 pN tension of human T-cells on CD19-only tension probes after 24 hr treatment with UCM, SCM, or ACM. Scale bar = 10 µm. Graph represents means ± SD and data points represent fluorescence intensity measurements of individual cells. Statistics were calculated using a one-way ANOVA with a multiple comparison test. UCM v. SCM *P* = 0.5903, UCM v. ACM *P* = 0.7449, SCM v. ACM *P* = 0.9526. *n* = 27 cells for UCM, 21 cells for SCM, and 27 cells for ACM.

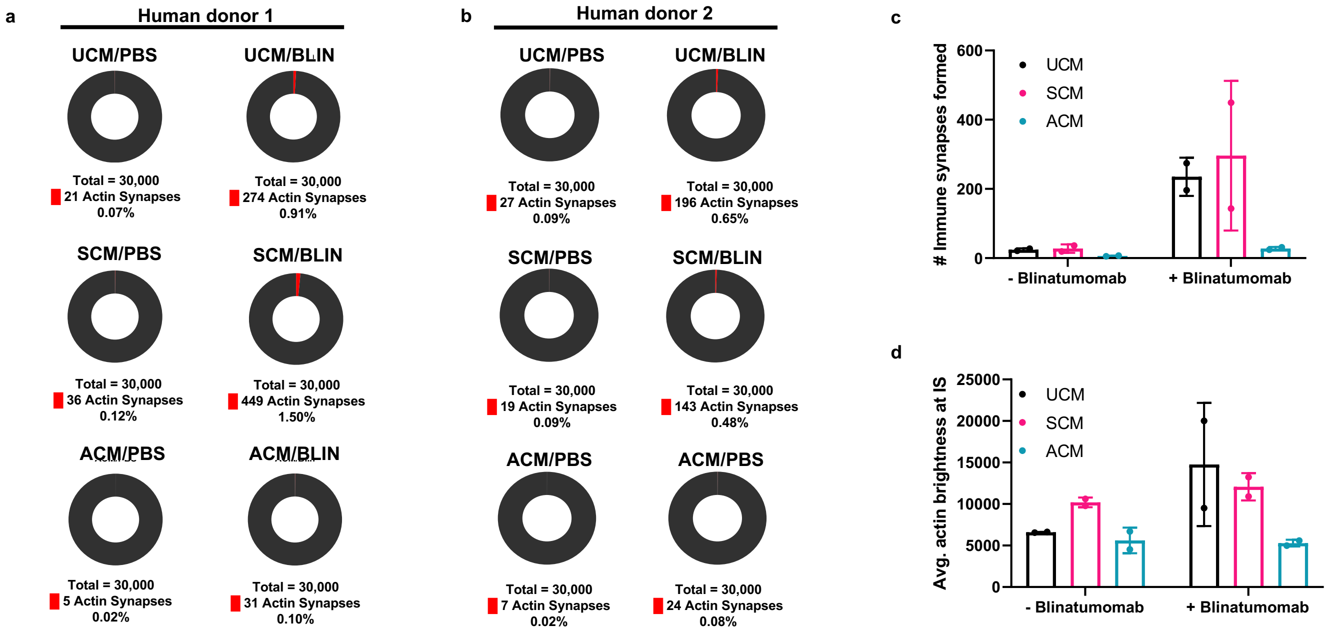


**Extended Data Fig. 9 | Additional quantification of blinatumomab-mediated immune synapses formed and actin brightness at the immune synapse in T-cell-tumor cell conjugates. a**, Pie charts showing the number of blinatumomab-mediated actin immune synapses formed in a co-culture experiment containing CD3+ T cells and Nalm6 B-cells in human donor 1 and **b**, human donor 2 performed in UCM, SCM, or ACM. **c**, Graphical representation of the number of blinatumomab-mediated immune synapses formed from both donors. **d**, Quantification of the average actin brightness at the immune synapse in co-culture experiments performed in UCM, SCM, or ACM. All data was acquired using the ImageStream cytometer.

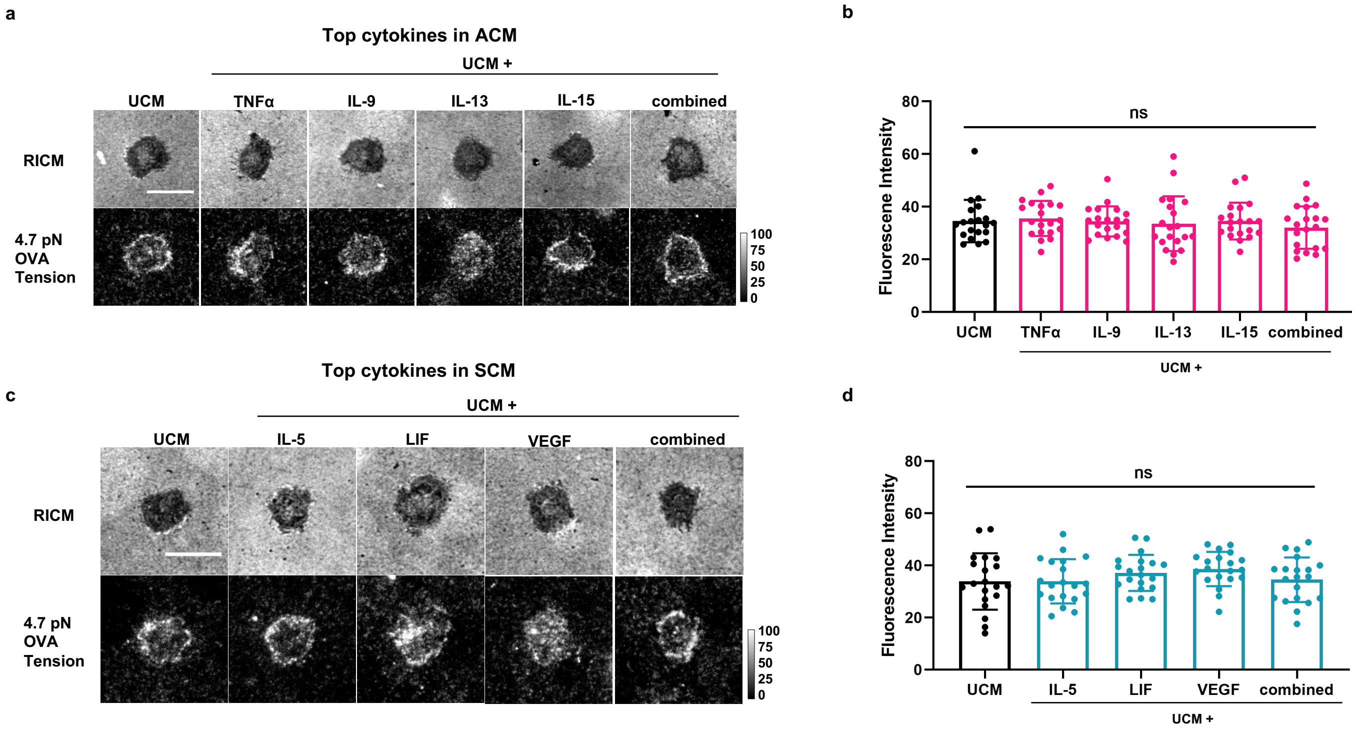


**Extended Data Fig 10 | Individual cytokines secreted by bone marrow stromal cells and adipocytes do not alter TCR biomechanics.** OT-1 T-cells were pre-treated for 24 hr with individual cytokines or a combination of the cytokines shown before being seeded on 4.7 pN hairpin probes presenting the OVA-pMHC. **a**, Representative images and **b**, quantification of 4.7 pN tension signal after treatment with top cytokines in ACM. $P$ = 0.8069. **c**, Representative images and **d**, quantification of 4.7 pN tension signal after treatment with top cytokines in SCM. $P$ = 0.2744. Scale bar = 10 µm. Graphs represent means ± SD and data points represent fluorescence intensity measurements of individual cells. Statistics were calculated using a one-way ANOVA. $n$ = 1 animal, 20 cells per condition.

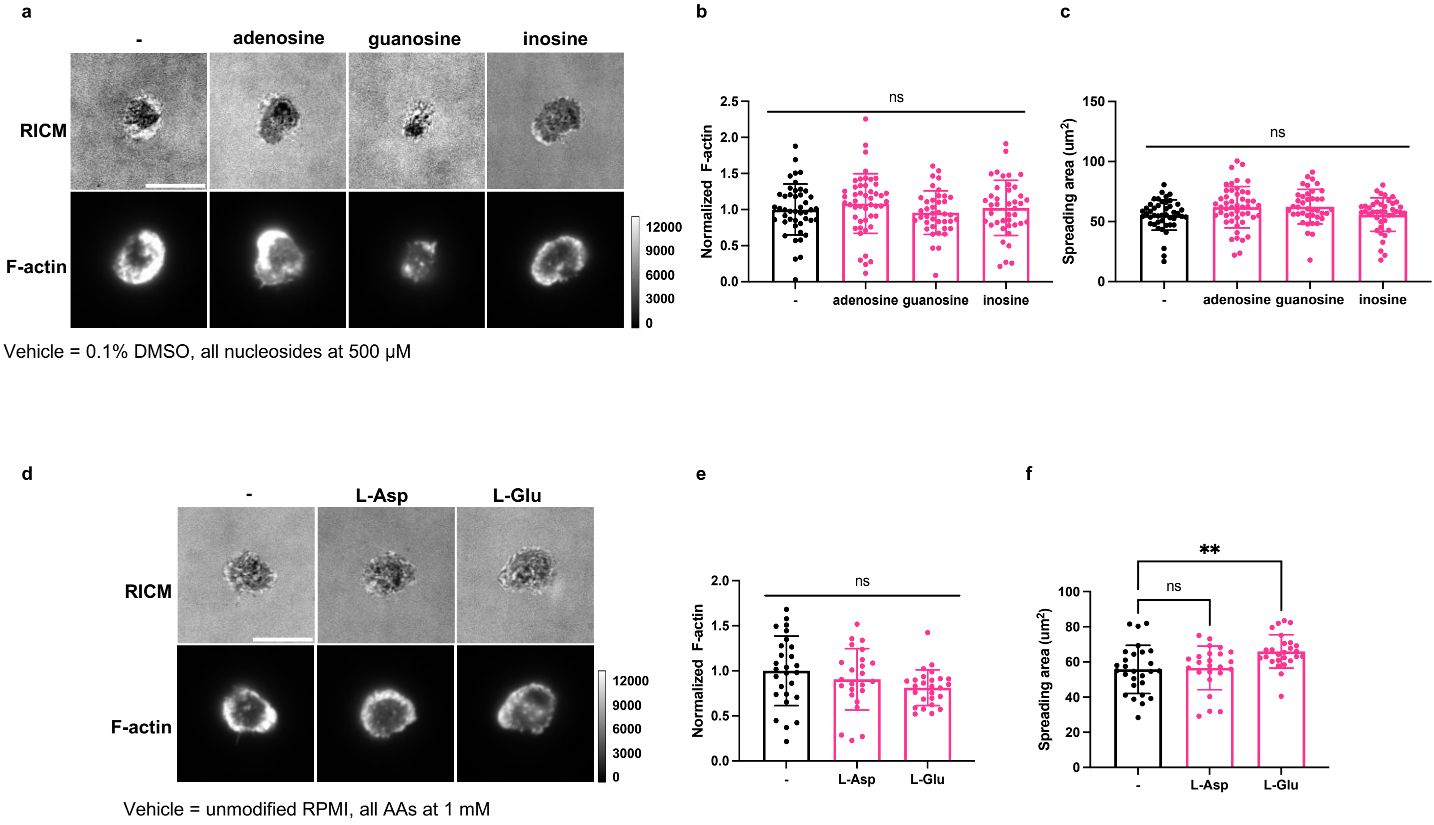


**Extended Data Fig 11 | Individual metabolites secreted by adipocytes do not alter F-actin polymerization or T cell adhesion. a**, Representative images of actin polymerization (F-actin) in OT-1 T-cells treated with nucleoside metabolites for 24 h and **b**, quantification of F-actin intensity and **c**, cell spreading area. N = 3 mice. **d**, Representative images of OT-1 T cells treated with amino acid metabolites for 24 h and **e,** quantification of F-actin intensity and **f**, cell spreading area. **P = 0.0052. N = 20-30 cells. Statistics were calculated using a one way ANOVA with a Tukey's multiple comparisons test.

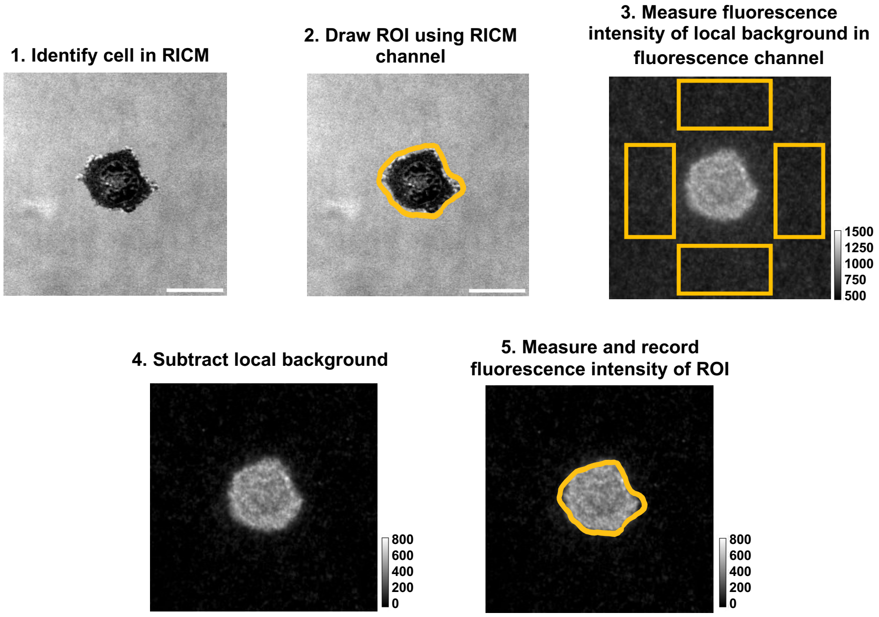


**Extended Data Fig. 12 | Workflow for measuring the fluorescence intensity of tension probe images and fluorescent staining in FIJI (ImageJ).** Adhered T-cells are identified in the RICM channel. The RICM channel is used to draw a region of interest (ROI) around each cell and each ROI is saved to the ROI manager. Next, the local background intensity is measured in the fluorescence channel (DAPI, TRITC, or CY5) by drawing 4 rectangular regions around the cell and measuring the mean intensity in each of those regions. The mean background fluorescence is averaged from the 4 regions and then subtracted from the image. The fluorescence intensity of the cell is then measured and recorded using the ROI drawn in step 2. Scale bar = 10 μm.

Western blot source data:
GAPDH

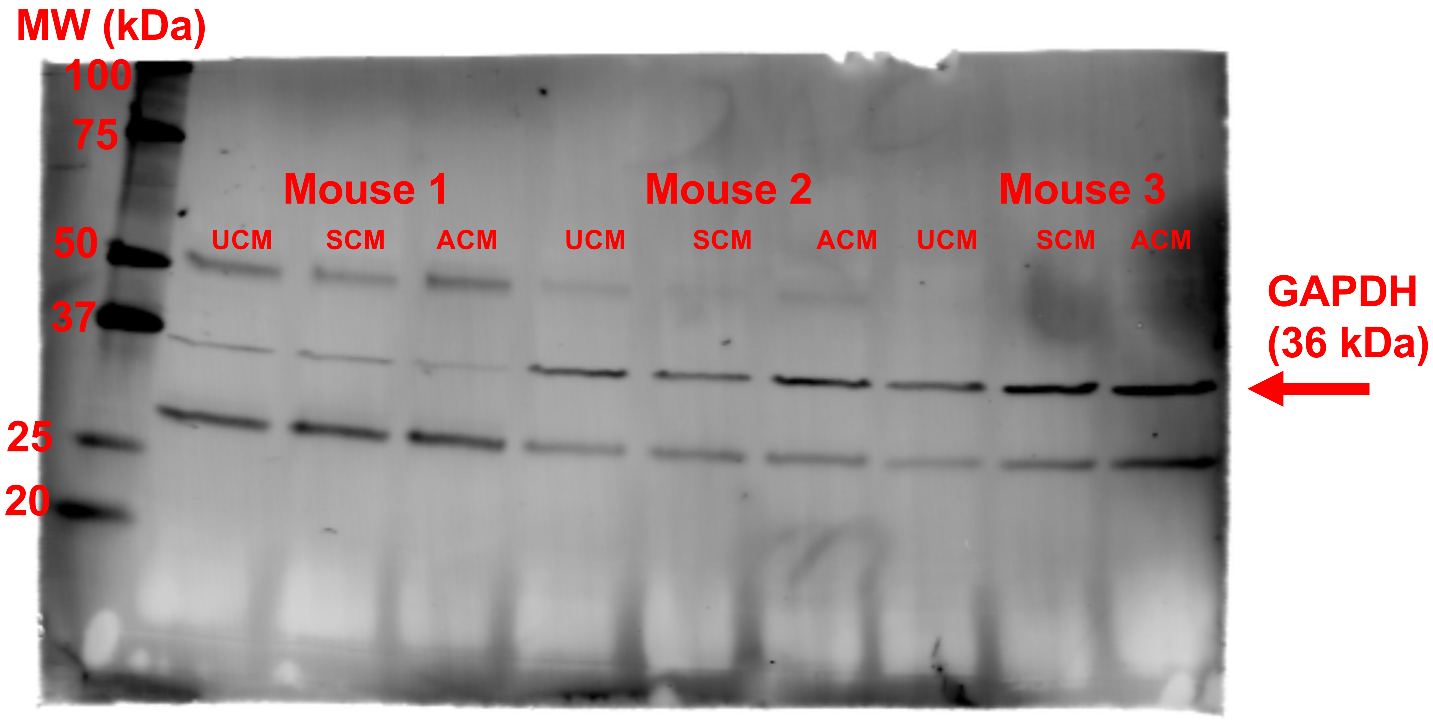

Western blot source data: β-actin

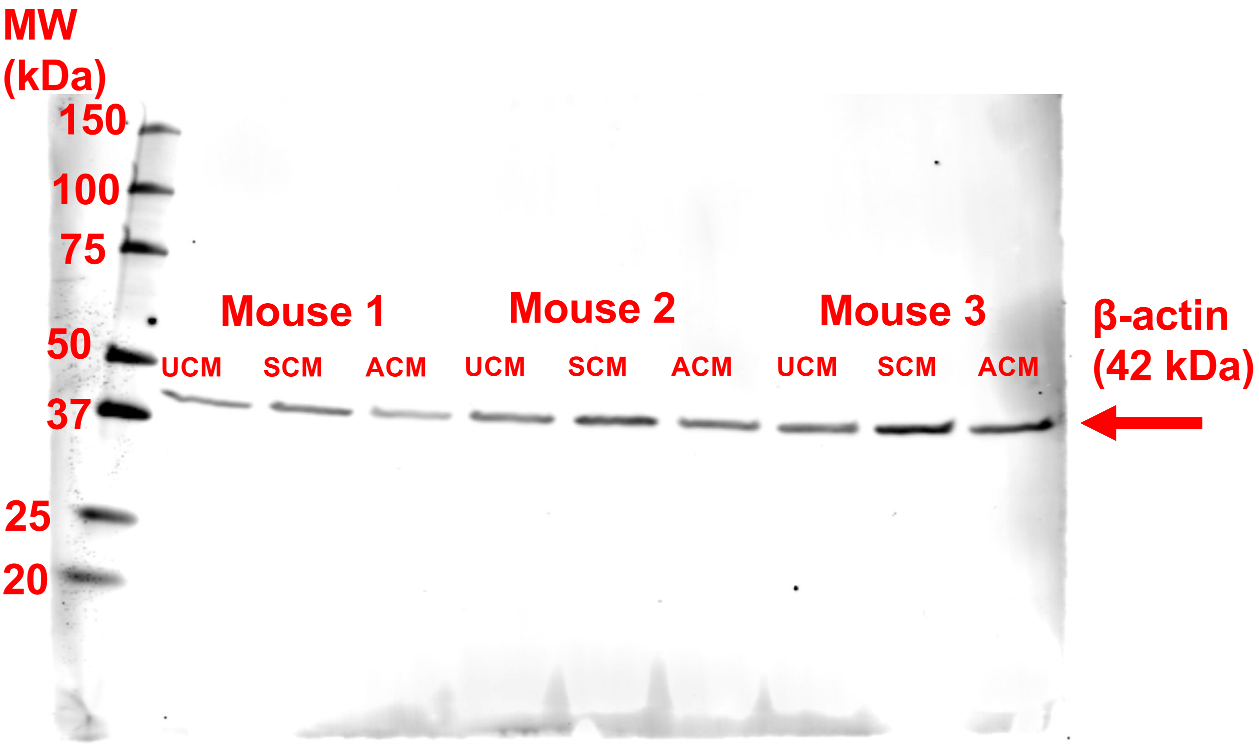
